# Biologically grounded locality priors close the data gap for vision transformers in neural prediction

**DOI:** 10.64898/2026.09.03.749141

**Authors:** Matteo Farina, Pietro Zamberlan, Arno Onken, Ulisse Ferrari

## Abstract

In computer vision, transformers have largely replaced convolutional networks, but not in neuroscience: vision transformers predict mouse V1 responses well only when trained on thousands of neurons. With no built-in locality, they must learn from data where each neuron looks. Electrophysiological experiments now commonly record from tens of neurons, with a few thousand stimuli, two orders of magnitude below the scale at which transformers are typically trained; with this little data, a transformer that sees the whole image tends to overfit, and convolutional neural networks and Gaussian processes perform better. We address this problem in the retina by restricting what the transformer sees: for each neuron, a differentiable crop keeps only the image features inside a small circle centered on its receptive field, *O*(*r*^2^) tokens for a radius *r* rather than the whole image, and a transformer, shared by all neurons, processes them. We call this model circular receptive-field vision transformer (CiRF-ViT) and test it on multi-electrode-array recordings of 41 mouse and 49 salamander retinal ganglion cells. Against CNN and Gaussian-process baselines, and a plain vision transformer with the same core that sees the whole image, CiRF-ViT reaches the highest mean noise-corrected correlation on both datasets (0.89 on mouse, 0.93 on salamander). We also tested the models with the local spike-triggered average (LSTA), the measured change of a cell’s response to small changes of a natural image, which for a model is the gradient of its response with respect to the image. For some cells, the LSTA shows a polarity inversion: around some images the cell responds like an ON cell and around others like an OFF cell. CiRF-ViT, never trained on this measure, reproduces this polarity inversion, which a linear-nonlinear model cannot capture by construction, and agrees with the measured LSTA as well as the baselines do. A biologically grounded, per-neuron locality prior is therefore enough to make transformers competitive for retinal response prediction well below their usual data scale.

## 1 Introduction

Deep encoding models of visual neurons increasingly rely on architectures imported from computer vision, whose gains are typically demonstrated at computer-vision scale: thousands of simultaneously recorded neurons and hundreds of thousands of stimulus presentations. Most electrophysiology does not look like this. A multi-electrode-array recording from an isolated retina, an acute cortical preparation, or a session with a behaving animal typically yields tens of neurons and a few thousand stimulus–response pairs, two to three orders of magnitude less. An architecture that performs best with large datasets does not necessarily do so when data are scarce. Here we ask which architecture is best suited to neural response prediction when both the number of neurons and the number of stimuli are small, and we study this question in the retina, where a receptive-field map of every recorded neuron is part of most recording protocols.

One family of encoding models fits each neuron independently, with a hand-designed inductive bias standing in for the missing data. Linear–nonlinear (LN) models project the stimulus onto a single spatial filter followed by a static nonlinearity; Gaussian-process (GP) models replace the filter with a kernel whose locality and smoothness are learnable priors (Goldin et al., 2023). Encoding locality directly in the model keeps them sample-efficient, but caps their capacity: a single filter or a smooth kernel cannot represent higher-order, nonlinear feature interactions, so their performance saturates early.

A second family shares a nonlinear feature extractor across the recorded population, and confines each neuron’s identity to a lightweight readout: the “what/where” separation of a convolutional core and a factorized spatial-and-feature readout (Klindt et al., 2017; Cadena et al., 2019; Lurz et al., 2021). Because the core is shared, and convolution is local and translation-equivariant, these models need far less data per neuron than an unconstrained network. Replacing the convolutional core with a vision transformer removes this built-in locality, since self-attention is global, in exchange for the ability to integrate long-range, contextual information. At sufficient scale the trade pays off: a shared transformer core with a per-neuron readout achieves state-of-the-art prediction of mouse V1 responses across sessions (Li et al., 2023). Below that scale it does not: in computer vision, plain vision transformers match convolutional networks only once training sets are large enough for locality to emerge from data (Dosovitskiy et al., 2021; Touvron et al., 2021).

This leaves a gap. Vision-transformer encoding models have only been demonstrated in the large-data regime that compensates for their lack of a locality prior (Li et al., 2023). Convolutional core-and-readout models exploit locality at the population level, but not at the level of the individual neuron: each cell’s receptive-field location is measured in a separate calibration in every one of these experiments, yet it is used only to weight the readout, never to determine what the shared feature extractor processes. No transformer-based encoding model uses this already-measured information to restrict what attention sees, although this prior could compensate for the small size of the dataset.

Here we supply this per-neuron locality prior: we ground a vision transformer’s attention in each neuron’s measured receptive field, and test on recordings of retinal ganglion cells whether this makes transformers competitive at a data scale where they are not expected to be. We develop and evaluate the method on the retina. Our contributions are the following.

- CiRF-ViT (circular receptive-field vision transformer), a vision transformer with a differentiable perneuron circular crop. The crop restricts the tokens seen by a shared transformer to a region of the feature map centered on the neuron’s receptive field, initialized from its spike-triggered average and refined during training. This reduces the number of attended tokens from *O*(*H* × *W*), for a feature map of height *H* and width *W*, to *O*(*r*^2^) for a crop radius *r*, and the benefit of this restriction is in how well the model learns from few data. With variable-length attention, the crop would also lower the cost of self-attention per neuron from *O*((*HW*)^2^) to *O*(*r*^4^).
- An evaluation on two multi-electrode-array retinal datasets (41 mouse and 49 salamander ganglion cells, a few thousand stimulus–response pairs each). CiRF-ViT reaches the highest mean noise-corrected correlation against LN, GP and CNN baselines and against a plain vision transformer with the same core but no crop (0.89 mouse, 0.93 salamander), with its largest advantage on the neurons the CNN predicts worst.
- Evidence that this is a data-efficiency effect: with the training set reduced to 10–20% of its size, CiRF-ViT remains competitive with a GP baseline built for the low-data regime.
- A zero-shot functional test, the local spike-triggered average (LSTA). CiRF-ViT reproduces the context-dependent polarity inversion that a linear model cannot capture and, within the band the experiment resolves, agrees with the measured LSTA as well as the CNN and GP baselines.

## 2 Related work

### Neural encoding models

Per-neuron models, such as LN models and Gaussian processes with a receptive-field-structured kernel (Goldin et al., 2023), fit each cell independently. Population models share a nonlinear feature extractor across neurons and confine per-cell identity to a factorized readout: the convolutional core-and-readout architecture of Klindt et al. (2017) has been scaled to macaque V1 (Cadena et al., 2019) and shown to generalize across recording sessions (Lurz et al., 2021). In all of these models, a neuron’s receptive-field location is used only at the readout, never to constrain which input regions the shared feature extractor processes. Li et al. (2023) replaced the convolutional core with a vision transformer and reported state-of-the-art prediction of mouse V1 responses, at the scale of thousands of neurons and hundreds of thousands of trials.

### Locality priors for data-efficient vision transformers

Outside neuroscience, several works give vision transformers part of the inductive bias of convolutional networks, locality and translation equivariance, while keeping global attention. Multi-head self-attention with a suitable relative positional encoding can express any convolution (Cordonnier et al., 2020). ConViT initializes each attention head as a convolutional kernel and lets a learnable gate relax it toward global attention where the data support it, matching convolutional networks at smaller training-set sizes (d’Ascoli et al., 2021). Swin restricts attention to local windows within a convolution-like hierarchy, with shifted windows restoring connectivity between them (Liu et al., 2021). ViT-P adds a learnable bias that suppresses attention outside a window at initialization and relaxes during training, closing much of the gap to convolutional networks on small datasets such as CIFAR-100 (Chen et al., 2022). Replacing the patch embedding with a few convolutions also stabilizes optimization at moderate data scale (Xiao et al., 2021). In all of these, the locality prior is the same for every image location, fixed by design or learned from the training distribution, and refers to no unit-specific information. In visual system identification such information is available, since every neuron’s receptive field is measured; CiRF-ViT grounds the locality prior in this measurement (Section 4).

## 3 Data

We use multi-electrode array (MEA) recordings of retinal ganglion cells (RGCs) collected by Goldin et al. (2022): a mouse dataset of 41 RGCs from a single retinal preparation, and a salamander (axolotl) dataset of 49 RGCs pooled from two preparations. The stimuli are natural images from the van Hateren dataset, at fixed mean luminance and RMS contrast, each flashed for 300 ms and interleaved with a uniform gray screen of equal duration (Figure 1a). The quantity to be predicted is each cell’s spike rate between 30 ms and 350 ms after image onset.

**Figure 1:**
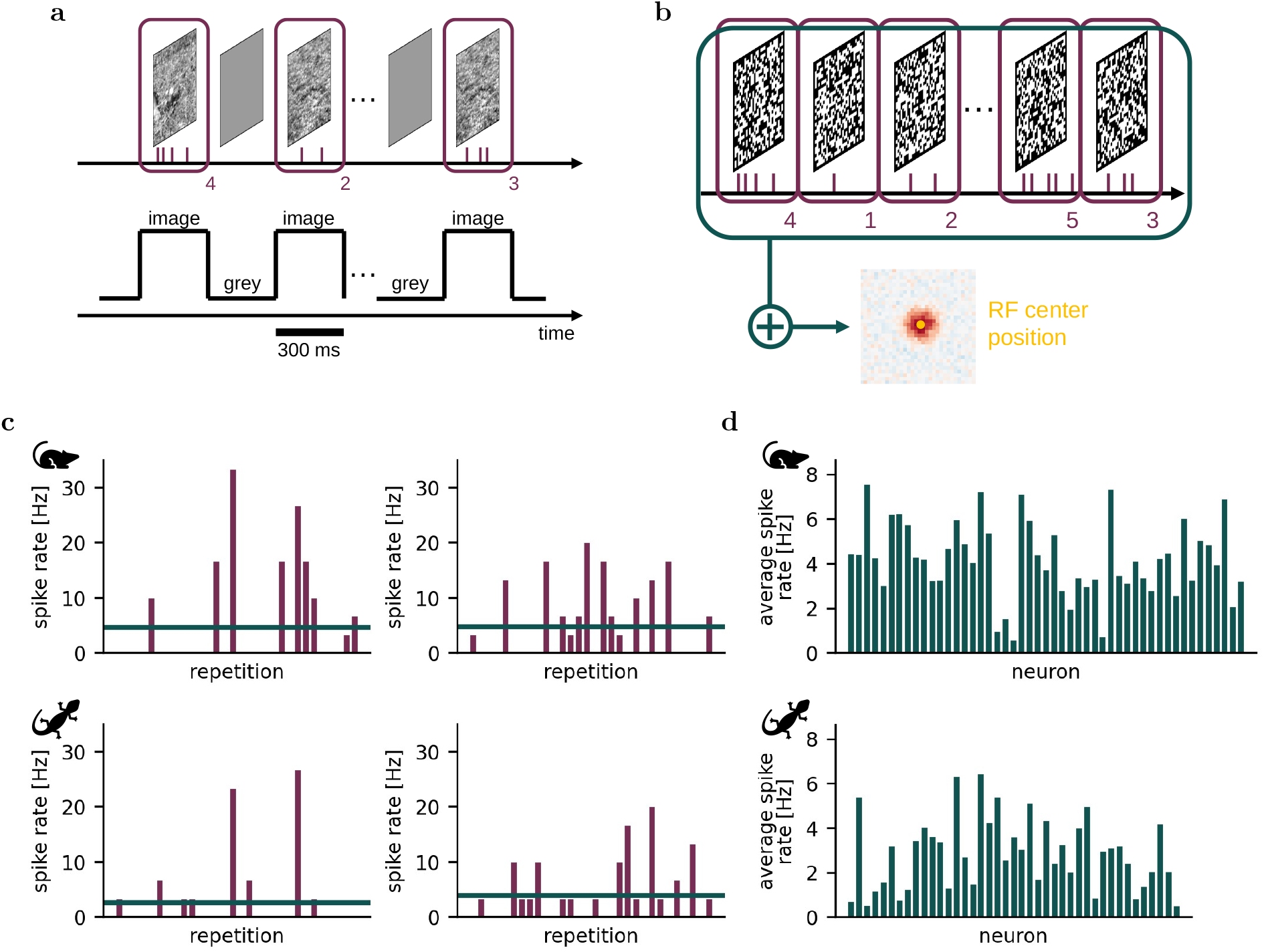
Dataset structure and response statistics. (a) Stimulus protocol. Natural images were presented for 300 ms each, separated by 300 ms of uniform gray. Spikes emitted during each image presentation were counted per neuron, yielding one response vector per stimulus. (b) Receptive field estimation. Binary white-noise checkerboard stimuli were presented and spike-triggered averages computed per neuron. The RF center is taken as the location of the peak absolute value of the STA (right, example neuron). (c) Test-set response variability. Spike rates across the repetitions of two test-set images (30 repetitions for mouse, 20 for salamander), for one example mouse RGC (top) and one example salamander RGC (bottom). Horizontal lines indicate the mean rate across repetitions. (d) Mean spike rate over the full dataset for every neuron, mouse (top, *N* = 41) and salamander (bottom, *N* = 49).

The receptive field (RF) of each neuron, the region of the visual field that drives its responses, is localized with the spike-triggered average (STA). In a separate stimulation block, binary white-noise checkerboards are presented at 30 Hz; the STA is the average of the frames that preceded a spike, weighted by the number of spikes each elicited, that is, the cross-correlation between stimulus and spike train, a first-order linear estimate of the neuron’s selectivity. We take the RF center as the location of the peak absolute value of the STA summed over time (Figure 1b); these centers initialize the locality prior of Section 4.

Throughout the paper, *receptive field* refers only to this physiological quantity, whose center we estimate from the STA and whose profile is approximately Gaussian. Two other spatial extents appear in the model and are named differently. The *crop window* of a neuron is the disk of radius *r*_*n*_, on the model’s feature map, that the model processes for that neuron (Section 4.2): a hyperparameter, not a measurement. The *token footprint* is the region of the input image that a single token depends on, set by the kernel size and stride of the tokenizer; like the theoretical receptive field of any convolutional unit it is a square, while its effective receptive field is narrower and roughly Gaussian (Luo et al., 2016; Kim et al., 2023).

All images are presented to the models at 108 × 108 pixels. The development set consists of 3160 images for both datasets, each presented once. The test set consists of 30 held-out images, repeated 30 times on mouse and 20 times on salamander; the response to each test image is the spike rate averaged over repetitions, since responses to the same image fluctuate considerably (Figure 1c). Mean firing rates range from below 1 Hz to about 8 Hz (Figure 1d). With a few thousand stimulus–response pairs for tens of neurons, both datasets are about two orders of magnitude below the scale at which vision transformers are typically trained for neural prediction.

## 4 Architecture

### 4.1 Benchmark models

**LN model** is a per-cell linear–nonlinear model, fitted independently to each neuron. Each input image is cropped to a 64 × 64 window centered on the neuron’s receptive-field center, and projected onto a single linear filter constrained to unit norm (MaxNorm, ∥*w*∥ ≤ 1). The linear response is passed through a parametric softplus nonlinearity, with a learnable output scale *α*, input gain *β*, and threshold *λ*. These are initialized so that at zero drive the output matches the neuron’s mean firing rate, which guarantees a strictly positive predicted spike rate. The model is trained with a Poisson negative log-likelihood loss, plus an L1 sparsity penalty and a squared-gradient smoothness penalty on the filter.

**GP model** fits an independent sparse variational Gaussian process to each neuron, following Goldin et al. (2023). Images center-cropped to 64 × 64 enter through an arc-cosine kernel (Cho & Saul, 2009), with learnable receptive-field position and size, defining a Gaussian envelope over pixels centered on the neuron’s RF center, and a smoothness parameter that sets the finest spatial scale the response can depend on; the kernel defines the prior over a latent function *λ*(*x*) of the image *x*. Spike counts follow a Poisson likelihood with rate exp(*A* · *λ*(*x*) + *λ*_0_), with learnable gain *A* and baseline *λ*_0_. Inference uses 300 inducing points and maximizes the ELBO, alternating variational, hyperparameter and likelihood-parameter updates. The same hyperparameters are used for both datasets (Table 3; see also Figure 2a).

**Figure 2:**
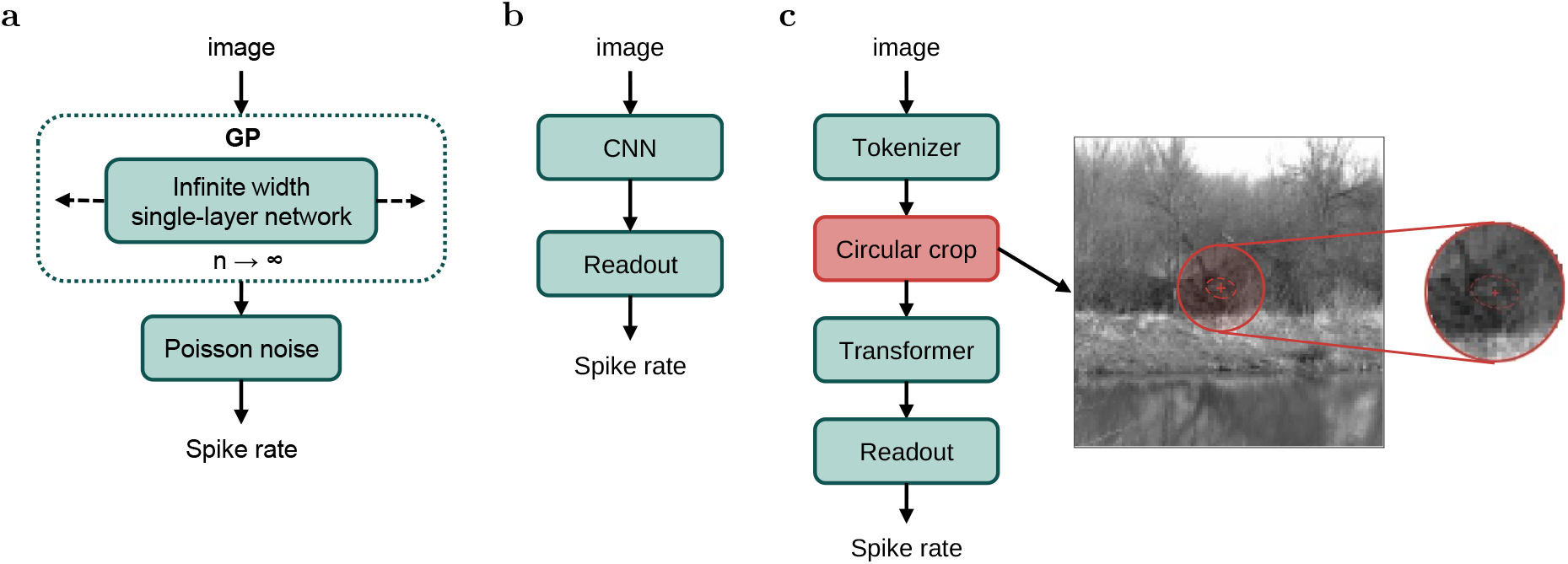
Model architectures. (a) Gaussian-process (GP) baseline. (b) CNN baseline: a shared convolutional core followed by a per-neuron factorized readout. (c) CiRF-ViT: a shared CNN tokenizer, a per-neuron differentiable circular crop centered on the neuron’s receptive field, shared transformer blocks, and a perneuron factorized readout.

**CNN model** consists of a convolutional core, with two convolutional layers shared across all neurons, and a per-neuron factorized readout (Klindt et al., 2017) that maps the features to the predicted spike rate. The readout applies a non-negative spatial mask *u* to the feature map, combines it with non-negative per-channel feature weights *v*, and passes the result through a softplus activation. The model is trained with a Poisson negative log-likelihood loss. Three regularization terms are used: a Laplacian smoothness penalty on the convolutional kernels, a distance-weighted spatial penalty on *u*, and an L1 sparsity penalty on the feature weights *v*. Detailed hyperparameters are reported in Table 2; see also Figure 2b.

### 4.2 CiRF-ViT architecture

CiRF-ViT consists of a shared tokenizer, a differentiable per-neuron crop, shared transformer blocks, and a per-neuron readout. We call *tokenizer* the module that turns the input image into the tokens processed by the transformer. Our tokenizer is a single convolutional layer (convolution, batch normalization and GELU), shared across all neurons. A token is one spatial position of its output feature map, a feature vector computed from an overlapping neighbourhood of the image. For each (image, neuron) pair, the crop extracts the features inside a circle centered on a point initialized at the neuron’s receptive-field center; bilinear sampling makes these center coordinates trainable by backpropagation (Section A.4). The circle is the same for all neurons, with the largest of the per-neuron radii; a per-neuron binary mask then excludes the features outside the neuron’s own radius, a fixed hyperparameter. The mask is applied both in the attention, as a key-padding mask, and in the readout’s spatial softmax. The cropped token sequences of all neurons are processed by shared transformer blocks, and a factorized readout maps them to the predicted spike rate through a per-neuron softmax-weighted spatial pooling, per-channel feature weights, and an ELU + 1 activation that keeps the output positive, as the Poisson loss requires. Models are trained with the Poisson negative log-likelihood and Adam; the per-neuron radii are selected by Bayesian hyperparameter search, jointly with the other hyperparameters (Table 1; see also Figure 2c).

**Table 1:** Hyperparameters of CiRF-ViT (transformer + per-neuron crop).

|  | **Mouse** | **Salamander** |
|---|---|---|
| Tokenizer channels / layers | 32 / 1 | 32 / 1 |
| Tokenizer kernel size | 3 | 11 |
| Tokenizer stride | 2 | 2 |
| Dropout | 0.1 | 0.1 |
| Transformer blocks / heads | 2 / 1 | 2 / 1 |
| MLP hidden dimension | 64 | 64 |
| Per-neuron radii (image px; effective: ×0.981) | 4–25 (mean 13.1) | 4–25 (mean 15.5) |
| Parameters | 38,956 | 46,732 |
| Batch size | 32 | 32 |
| Optimizer / learning rate | Adam / 10<sup>−3</sup> | Adam / 10<sup>−3</sup> |
| Max epochs / early-stop patience | 200 / 30 | 200 / 30 |
| Loss | Poisson NLL | Poisson NLL |

**Table 2:**
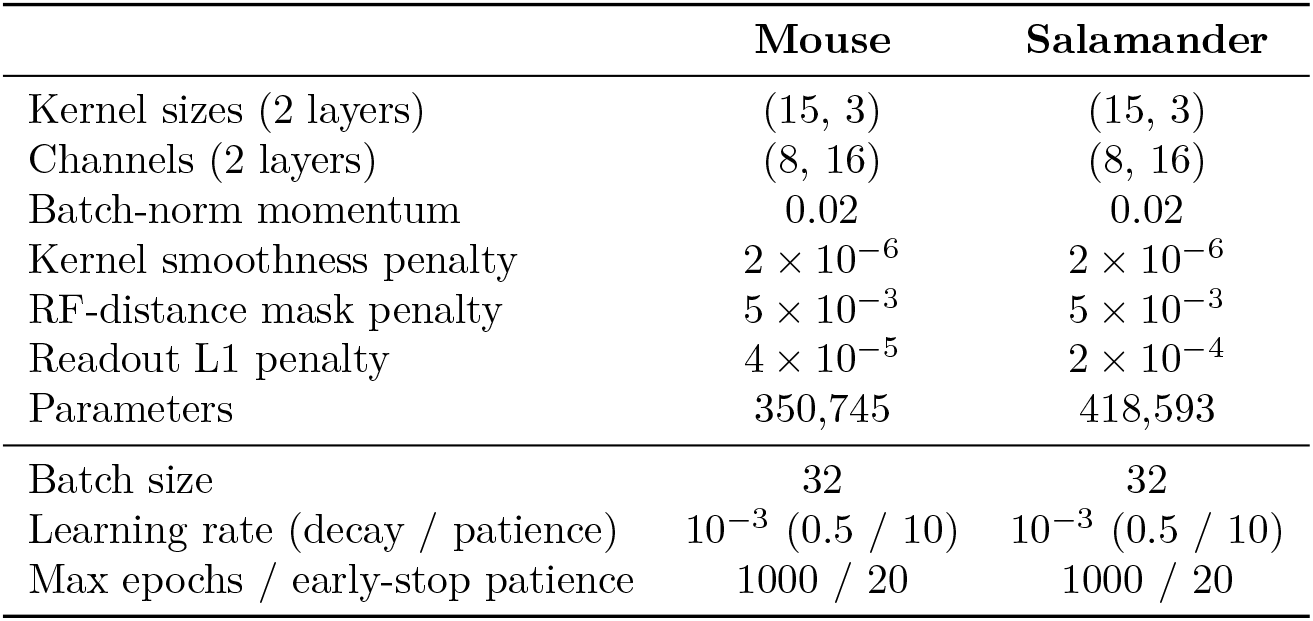
Hyperparameters of the CNN baseline (shared convolutional core + factorized readout).

**Table 3:**
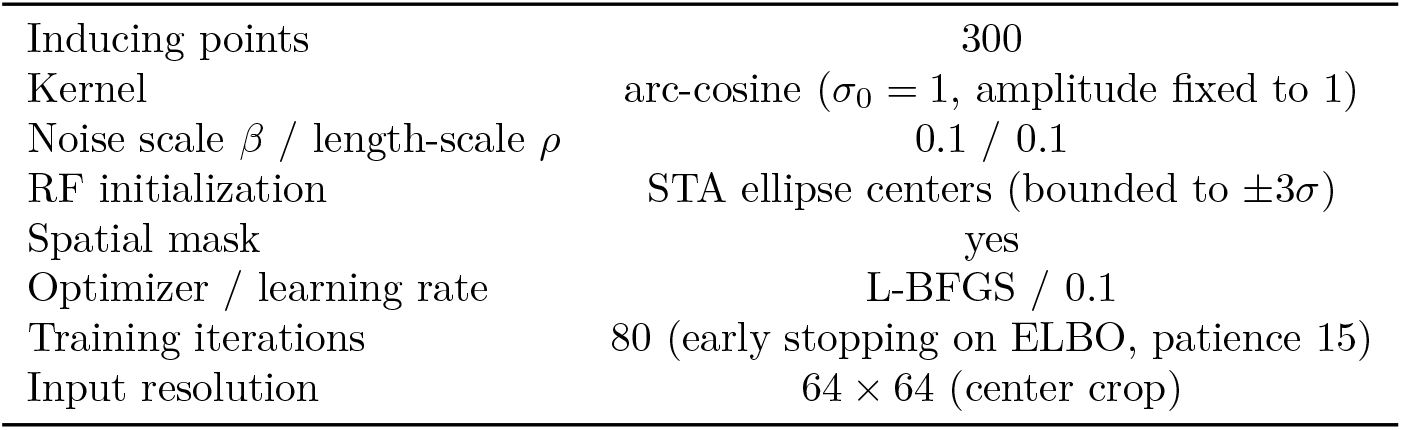
Hyperparameters of the GP baseline (Goldin et al., 2023), identical for both datasets.

|  |  |
|---|---|
| Inducing points | 300 |
| Kernel | arc-cosine (σ<sub>0</sub> = 1, amplitude fixed to 1) |
| Noise scale β / length-scale ρ | 0.1 / 0.1 |
| RF initialization | STA ellipse centers (bounded to ±3σ) |
| Spatial mask | yes |
| Optimizer / learning rate | L-BFGS / 0.1 |
| Training iterations | 80 (early stopping on ELBO, patience 15) |
| Input resolution | 64 × 64 (center crop) |

## 5 Benchmarks

All results are obtained with 5-fold cross-validation: the development set is split into five folds, the models are trained on four of them, with a validation split for early stopping and hyperparameter selection, and evaluated on the held-out test images (Section 3). Performance is the noise-corrected correlation (*CC*_norm_; Schoppe et al., 2016) between the predicted spike rate and the repeated test responses, computed per neuron: the correlation normalized by the highest value the trial-to-trial variability of the recording allows, so that an ideal model scores 1 (Section A.5).

Besides the LN, GP and CNN baselines, the comparison includes a plain vision transformer: CiRF-ViT without the crop, so that every neuron attends to, and is read out from, the whole feature map (the *no crop* variant of Section A.1). It isolates the contribution of the per-neuron locality prior from that of the architecture.

CiRF-ViT is compared with each reference model by a two-sided Wilcoxon signed-rank test (Wilcoxon, 1945) on the per-neuron values of the box plots (Figure 3a, Figure 4e), each averaged over the five cross-validation folds, paired neuron by neuron (*N* = 41 on mouse, *N* = 49 on salamander). No correction for multiple comparisons is applied, as each test addresses a distinct, pre-specified comparison.

**Figure 3:**
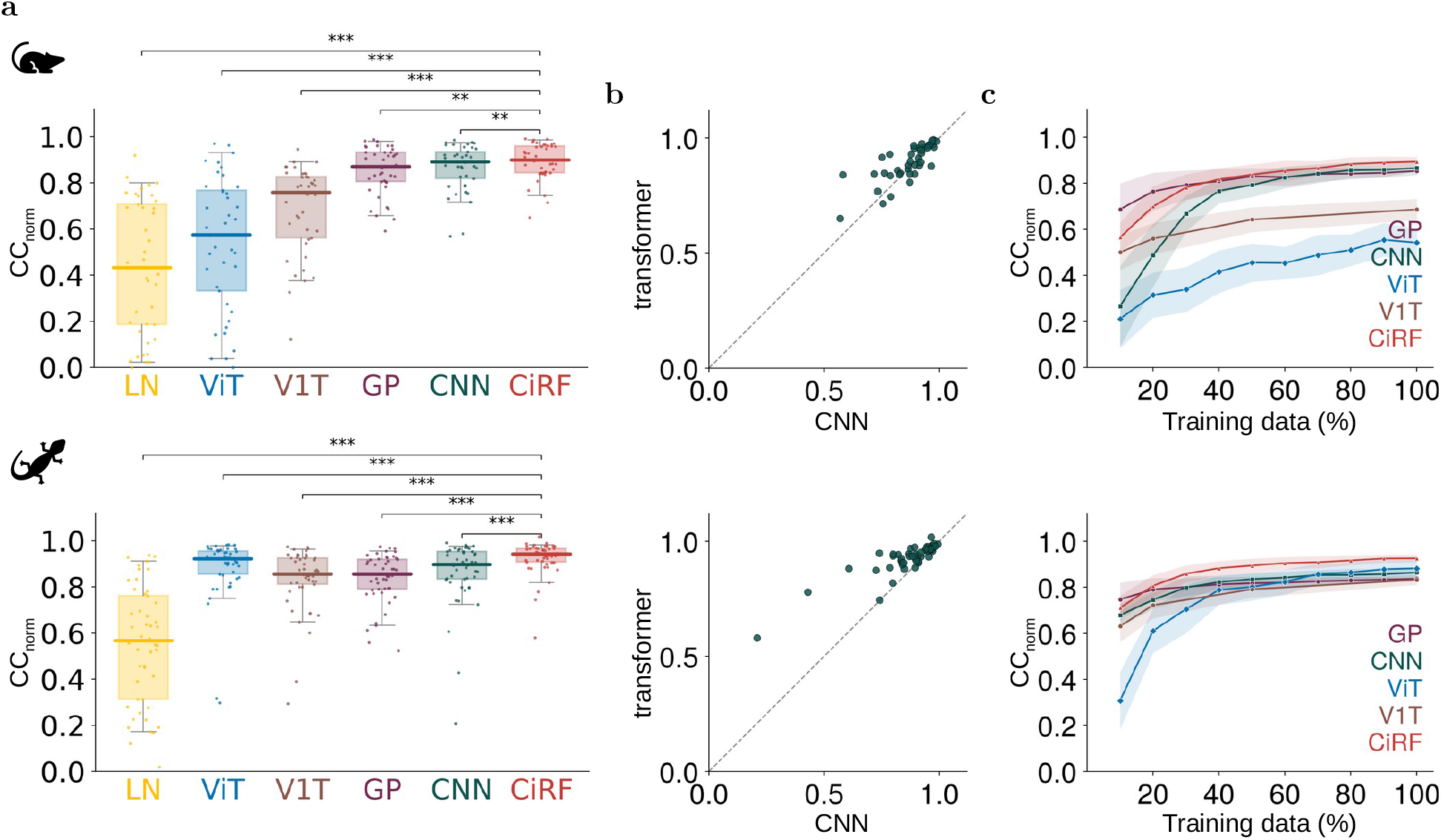
Prediction performance across models and datasets (top: mouse, *N* = 41 RGCs; bottom: salamander, *N* = 49 RGCs; 5-fold cross-validation). (a) Noise-corrected correlation (*CC*_norm_) on the test set for the LN, GP, CNN, plain ViT (CiRF-ViT without the crop; labeled ViT), V1T (the best pretrained core, fine-tuned; Section A.2) and CiRF-ViT (labeled CiRF) models, ordered from the lowest to the highest mean over the two datasets. Dots: neurons; boxes: interquartile range; thick line: median; whiskers: 5th to 95th percentiles. Stars: paired per-neuron Wilcoxon signed-rank test against CiRF-ViT (* *p <* 0.05, ** *p <* 0.01, *** *p <* 0.001). (b) Per-neuron *CC*_norm_ of CiRF-ViT (y-axis, labeled transformer) against the CNN (x-axis); dashed line: identity. (c) *CC*_norm_ as a function of the fraction of the training and validation images used; shaded regions: standard deviation across runs, computed per neuron and averaged over neurons (five runs per point; for the plain ViT and V1T, one per cross-validation fold; V1T at 10, 20, 50 and 100% of the data).

**Figure 4:**
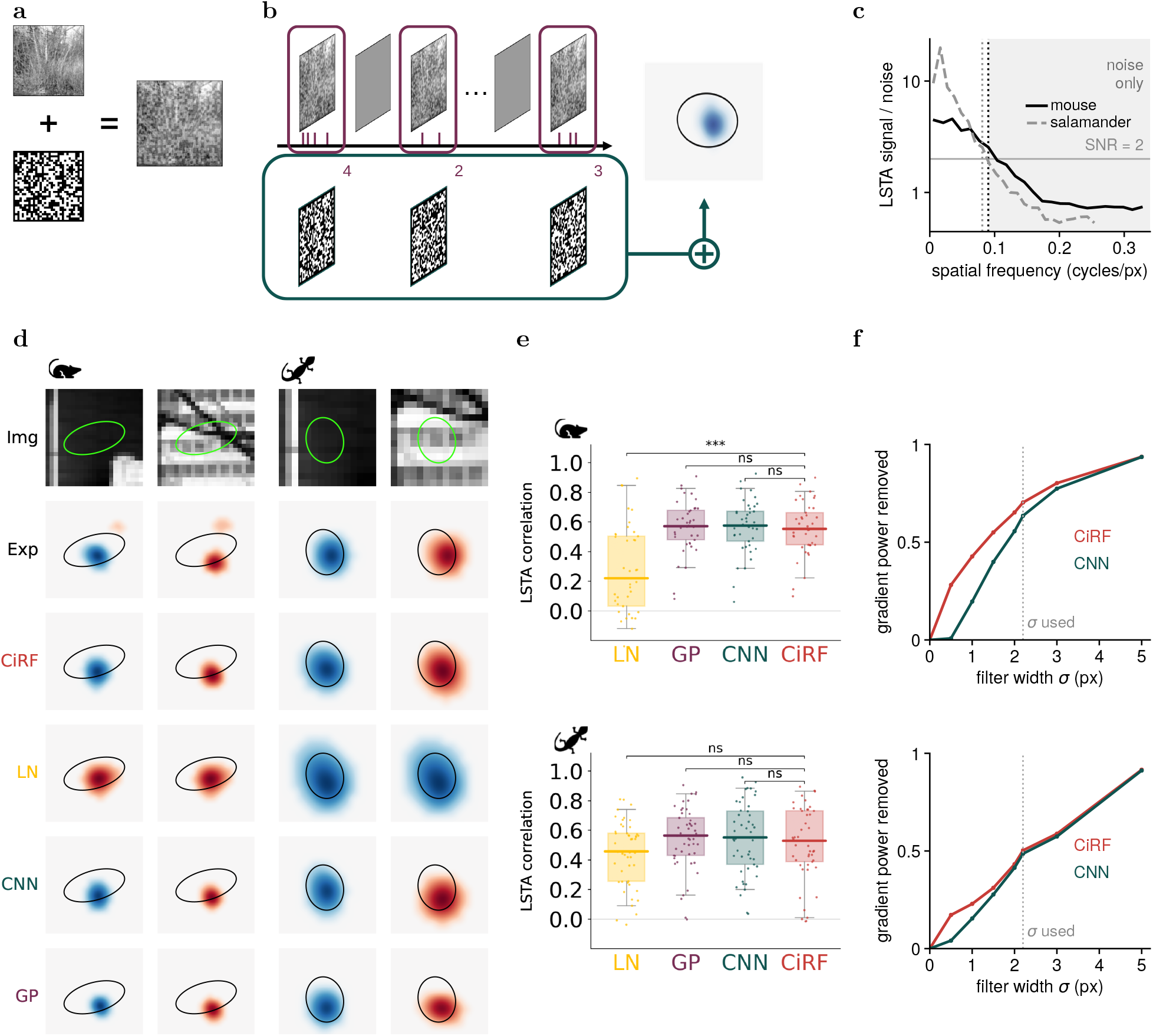
Local spike-triggered average (LSTA). (a) Stimulus: binary white-noise checkerboards, identical to those used for the STA (Figure 1b), superimposed at low contrast on natural images. (b) The LSTA is the spike-triggered average of the noise component of these stimuli. (c) Resolution of the experimental LSTA: ratio between its power inside the receptive-field window and in an identical window far from the receptive field, as a function of spatial frequency. Beyond the dotted lines (ratio *<* 2) the maps contain only estimation noise, and the comparison in (d, e) is low-passed accordingly (Section A.6). (d) Experimental LSTAs and model gradients for two example neurons (one mouse, one salamander), each for two test-set images (the same neurons and images as in Figure 1c), shown band-limited. Top row: stimulus, with the receptive-field outline; second row: experimental LSTA; remaining rows: gradient of the predicted spike rate with respect to the input image. (e) Per-neuron correlation between the band-limited experimental LSTA and model gradient, for the LN, GP, CNN and CiRF-ViT models, on mouse (top, *N* = 41) and salamander (bottom, *N* = 49); dots: neurons; brackets: two-sided Wilcoxon signed-rank test between CiRF-ViT and each model (*** *p <* 0.001; ns, not significant). (f) Fraction of input-gradient power removed by a Gaussian low-pass of width *σ*, for CiRF-ViT and the CNN (Table 8), mouse (top) and salamander (bottom); the dotted line marks the width used in (d, e).

### 5.1 Prediction performance

With the full training set, CiRF-ViT achieves the highest mean *CC*_norm_ on both datasets: 0.89 on mouse and 0.93 on salamander (Figure 3a). On mouse its advantage over the CNN, the strongest reference model (0.87), is small, and both clearly outperform the LN and GP baselines. On salamander the gap is larger: CiRF-ViT predicts better for the large majority of neurons, and most for those the CNN predicts poorly (Figure 3b), lifting the low-performing tail of the population rather than only the cells that were already well predicted. Removing the crop, with every other component unchanged, costs most on mouse, where the plain ViT falls to 0.54, below every model except the LN, and less on salamander (0.88, between the CNN and CiRF-ViT); on both datasets the difference is significant (*p <* 0.001).

With less training data (Figure 3c), CiRF-ViT degrades, as expected, but at 10–20% of the data (roughly 300–600 training images) it remains comparable to the GP, which is designed for the low-data regime (Goldin et al., 2023), and it overtakes all baselines well before the full training set is reached. With the per-neuron locality prior, a vision transformer thus no longer needs the data scale at which such models are typically trained. Without it, it does: on mouse the plain ViT stays far below every other model at all data fractions, and on salamander it approaches the CNN only with most of the training set.

An alternative to a locality prior, when data are scarce, is to start from a transformer pretrained on other data. The best pretrained core we tested is the V1T core of Li et al. (2023), trained on the responses of mouse V1 to natural images, which we fine-tuned with the readout of CiRF-ViT over the whole token grid. With the full training set it reaches 0.69 on mouse and 0.83 on salamander, below both CiRF-ViT and the CNN (Figure 3a; *p <* 0.001 against CiRF-ViT on both datasets). It stays below CiRF-ViT at every training-set size (Figure 3c), including 10% of the data, where pretraining should help most (0.50 against 0.56 on mouse, 0.63 against 0.71 on salamander). The other experiments with pretrained cores, ViT-S/16 models trained on ImageNet and frozen cores, are presented in Section A.2.

## 6 Beyond prediction: probing retinal computation

Prediction accuracy measures how well a model fits the data it was trained on, not whether it has captured the underlying computation. As an independent functional test, we compare the models with the local spike-triggered average (LSTA) (Goldin et al., 2022), measured by superimposing a binary white-noise checkerboard on a natural image at low contrast (Figure 4a) and spike-triggered averaging the noise component alone (Figure 4b). Because the noise is a small perturbation around a fixed image, the LSTA estimates the neuron’s first-order sensitivity around that image: it is the experimental counterpart of the gradient of a model’s predicted spike rate with respect to the input image, which we compute for each trained model on the same images. The models are never trained on LSTA data, so this is a zero-shot test. The LSTA was recorded in a separate block, with stimuli that never appear in the training data, and it is measured around held-out test images. Nor do the training data carry any signal about gradients: the loss compares predicted and recorded responses at the training images only, so nothing in it specifies how a response changes when an image is perturbed, and whatever a model captures of the LSTA it has inferred from the responses to natural images. Most of the models’ gradient energy lies within the span of the images, but a part lies outside it (Figure 5): the 3190 images of the mouse dataset span at most 3189 of the 108^2^ = 11664 pixel directions, and 16% of the CNN’s and 35% of the transformer’s gradient energy falls along directions in which no image of the dataset varies, which the stimuli cannot constrain at all.

**Figure 5:**
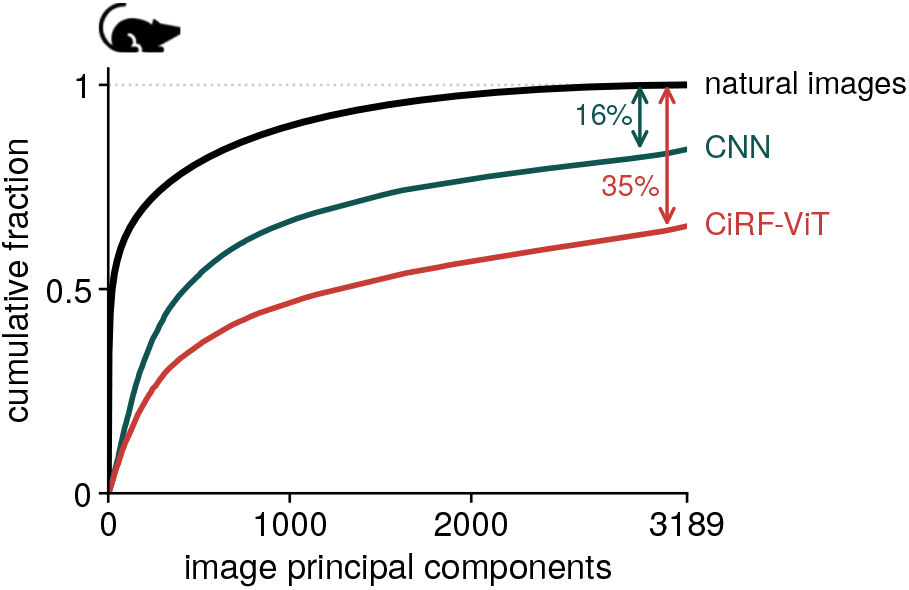
Gradient energy outside the span of the natural images (mouse dataset). Principal component analysis of all 3190 images of the dataset (training, validation and test, mean-subtracted) in the space of the 108^2^ = 11664 pixels: with 3190 images, at most 3189 components carry variance, and together they explain all of it by construction. Black: cumulative fraction of the image variance along these components. Colored: cumulative fraction of the energy of the input gradients of a CNN and CiRF-ViT (41 neurons, 8 LSTA images; squared projections summed over all gradients), normalized by their energy over all 11664 pixel directions. At the last component the image curve reaches 1, while the gradient curves stop at 0.84 and 0.65: the remaining 16% and 35% lie outside the span of the images.

The STA and the LSTA are related but answer different questions. The STA (Section 3) is measured without natural images and yields a single linear filter per neuron; mathematically, it is the LSTA computed around a uniform gray image. The LSTA extends it to natural-image contexts, where it can differ from the STA in shape and even in sign. Of the STA, CiRF-ViT receives only the location of its peak, two scalars per neuron that initialize the crop center; it receives neither the shape, the amplitude nor the polarity of the filter, nor its dependence on context, which it has to learn from the natural-image responses alone. Variants of the model that never receive the STA also agree with the LSTA, to the extent that they predict the responses well (Section A.1, Figure 11).

Depending on the image context, the LSTA can invert its polarity relative to the STA, switching a neuron from ON- to OFF-type selectivity (Goldin et al., 2022): a nonlinear effect that a linear model cannot reproduce by construction. CiRF-ViT reproduces the inversion whenever it is present experimentally (Figure 4d), while the LN model cannot.

The quantitative comparison has to respect what the experiment can measure. Estimated from a finite number of spikes, the LSTA carries signal only at coarse spatial scales: compared with an identical window far from the receptive field, where the map contains only estimation noise, its power is less than twice the noise for spatial periods finer than 11 px on mouse and 12 px on salamander (Figure 4c). In that band there is nothing for a model to agree with, but an unfiltered correlation would still penalize a model for its content there, which enters the model’s variance in the denominator of the correlation, and the more so the finer its gradient. We therefore low-pass the experimental LSTA and every model’s gradient with the same Gaussian filter (*σ* = 2.2 px, half-amplitude point at the measured cutoff) before correlating them (Section A.6). This changes the measurement, not the models.

Within this band, CiRF-ViT, the CNN and the GP agree with the measured LSTA equally well on both datasets (Figure 4e; Wilcoxon signed-rank, *p >* 0.05 for CiRF-ViT against the CNN and against the GP), while the LN model is worse on mouse (*p <* 0.001). The LSTA is not a second accuracy benchmark, on which the best predictor should also win, but a check that a model’s first-order sensitivity is consistent with the measured one: a model whose higher accuracy came from physiologically implausible sensitivities would fall significantly below the baselines, and CiRF-ViT does not. The band limit removes more from CiRF-ViT than from the CNN (Figure 4f): 70% of its gradient power on mouse and 50% on salamander, against 64% and 49%, and at the finest scales (*σ* = 0.5) 28% against less than 1% on mouse (Table 8). CiRF-ViT’s gradients are thus far richer in high-frequency components, exactly the content the experimental LSTA cannot resolve.

## 7 Discussion

We asked whether a vision transformer can be made competitive for predicting retinal responses two orders of magnitude below its usual data scale, and found, on both retinal datasets, that constraining its attention with information already available for every recorded neuron, the location of its receptive field, is enough. CiRF-ViT’s advantage over the convolutional baseline is concentrated on the neurons the CNN predicts worst, which suggests that the crop helps where a population-level locality prior is not specific enough to the cell, and it persists, in a weaker form, with a fraction of the training data. Within the band the experiment resolves, CiRF-ViT agrees with the measured LSTA as well as the CNN and GP; what the probe leaves open is the band it cannot resolve, where CiRF-ViT’s gradients carry far more content than the baselines’.

One reading of these results is that, for small-scale prediction of retinal responses, the trade-off between transformer and convolutional architectures is not fixed by data scale alone, but also depends on whether what the experimenter already knows is handed to the model as a prior or left to be rediscovered from data.

This does not establish that transformers are generally preferable; it shows that the expectation that they need scale regardless of what is known does not hold once a relevant, measured prior can be built into the architecture. Nor does pretraining remove the need for the prior: transformer cores pretrained on ImageNet or on mouse V1 responses, frozen or fine-tuned, stay below CiRF-ViT at every training-set size (Figure 3c and Figure 12). At this data scale, knowing where each neuron looks is worth more than the features these models transfer from other images or other neurons.

### Limitations

Our evaluation is confined to the retina. Although our study includes data recorded from two species (mouse and salamander) with different response statistics and receptive-field sizes, we have not tested whether the same prior is equally effective for encoding models of other visual areas or sensory modalities. Our model also requires a receptive-field center for every neuron. In the retina this is a mild requirement, since a white-noise calibration block is a standard part of most recording protocols; elsewhere receptive fields can be broader, less stable across a session, or not mapped. The center, however, need not come from a dedicated measurement: centers estimated from the natural-image training responses alone reproduce most of the result, and a coarse estimate loses little (Figure 9). Our findings indicate that providing an approximate center is sufficient, because training refines the center only locally (Table 6), and with random centers the model fails. The LSTA comparison, in turn, pertains only to the band the experiment resolves, where every model is smoothest. Finally, the crop radius is a per-neuron hyperparameter selected by Bayesian search, not learned end to end. The computational saving of the crop is also only theoretical: our implementation pads every neuron to the largest crop, much larger than the crop of most neurons, and runs dense attention once per neuron, so its attention cost is comparable to that of global attention (Section A.3); the benefit of the crop is in how well the model learns from few data, not in training time. The crop samples its tokens on a lattice of spacing (*W* − 1)*/W* ≈ 0.981 feature-map positions rather than 1, a convention of the implementation kept because the radii were selected with it (Section A.4).

Like any inductive bias, the per-neuron crop has a cost, and so does the convolutional bias it replaces. A convolutional core applies the same local kernel at every location: interactions between distant image regions can only be built up through depth, and the same features are computed for every neuron, wherever its receptive field lies. The crop makes the opposite trade: CiRF-ViT cannot respond to anything outside each neuron’s crop window, so stimulus structure beyond it, such as a far surround or a wide-field modulation, is invisible to the model, and a larger radius only partly compensates, since it moves the model back toward the unconstrained, data-hungry regime the crop is meant to avoid.

CiRF-ViT’s gradients are also far richer than the CNN’s at the finest scales (Figure 4f), and nothing in the architecture suppresses this content, unlike the smoothness priors of the LN and GP models. Whether it is a genuine feature of retinal processing or a predictive shortcut, these data cannot say: in that band, the measurement contains only noise.

Where the prior is enforced also matters. On mouse the crop works best inside the core, for the transformer as for a convolutional core with the same parameter count and interaction range, and no convolutional core reaches CiRF-ViT; on salamander restricting the crop to the readout, with global attention, is enough to match CiRF-ViT, and for the convolutional core the position of the crop makes no difference (Figure 6), showing that the best position of the crop depends on the dataset.

**Figure 6:**
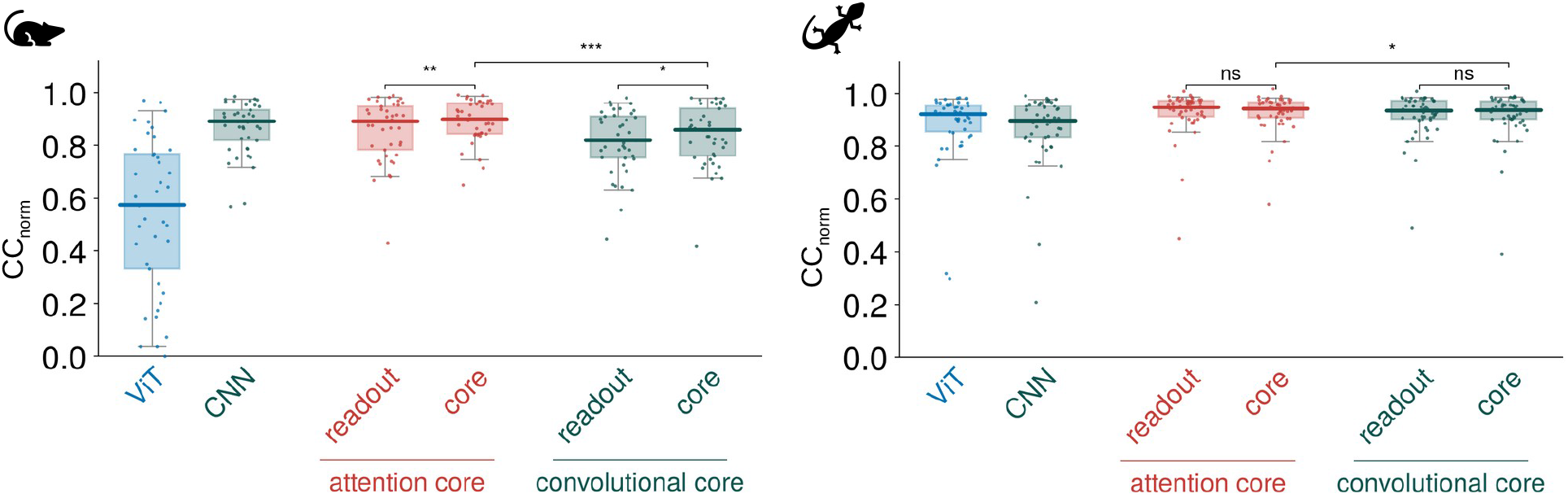
Attention against convolution, with the crop in the core or in the readout. Left: mouse; right: salamander. Per-neuron *CC*_norm_ on the test set, mean of five runs (one per cross-validation fold; for CiRF-ViT and the CNN, the five runs at the full training set of Figure 3). From left to right: the plain ViT and the CNN baseline of Figure 3; the transformer with the crop in the readout and in the core (CiRF-ViT); the convolutional core, matched to the transformer blocks in parameter count and interaction range, with the crop in the readout and in the core. Brackets: two-sided Wilcoxon signed-rank test, paired over neurons, between the two crop positions of each core and between the two cores with the crop in the core (* *p <* 0.05, ** *p <* 0.01, *** *p <* 0.001; ns, not significant). Dots: individual neurons. Mean and standard deviation of every combination in Table 4.

### Future directions

The high-frequency content of CiRF-ViT’s gradients is currently untestable: characterizing which image features it responds to would show whether the model has found something about retinal processing or a predictive shortcut, and a sensitivity measure built for noisy, high-dimensional settings could extend the model–data comparison into the band where a pointwise correlation with a noisy measurement is uninformative. Beyond the retina, applying the crop to a cortical dataset, with an independently measured receptive field per neuron, would show whether the low-data advantage is a property of retinal computation or of any encoding problem where each unit’s receptive field is known in advance.

**Table 4:** *CC*_norm_ of every combination of core and crop position (Figure 6), including the convolutional core without crop: mean ± standard deviation over the five runs. The CNN baseline restricts its readout with a soft distance penalty (soft mask) or, as a control, with the hard circular mask of CiRF-ViT (hard mask).

| Mouse (*N* = 41) |  |  | Salamander (*N* = 49) |  |  |
|---|---|---|---|---|---|
|  | **attention** | **convolution** |  | **attention** | **convolution** |
| no crop | 0.542 ± 0.020 | 0.616 ± 0.024 | no crop | 0.882 ± 0.013 | 0.881 ± 0.008 |
| crop in readout | 0.856 ± 0.004 | 0.812 ± 0.010 | crop in readout | 0.927 ± 0.003 | 0.918 ± 0.002 |
| crop in core | 0.893 ± 0.006 | 0.837 ± 0.012 | crop in core | 0.925 ± 0.002 | 0.917 ± 0.003 |
| CNN, soft mask | 0.866 ± 0.003 |  | CNN, soft mask | 0.864 ± 0.006 |  |
| CNN, hard mask | 0.876 ± 0.004 |  | CNN, hard mask | 0.856 ± 0.008 |  |

Taken together, these results suggest that, in the low-data regime, much of what a vision transformer would otherwise have to learn from data, namely where each neuron looks, can be supplied by a measurement the experimenter already has: when a neuron’s receptive field is known, restricting attention to it is enough to make the transformer competitive.

## Broader Impact Statement

This work uses only previously published recordings from isolated mouse and salamander retinas (Goldin et al., 2022); no new animal experiments were performed. For future data collection, its implication is a reduction: when each neuron’s receptive field is known, an accurate model can be trained from a few thousand stimulus presentations, and the receptive-field center it requires comes from a short white-noise block that is already part of most retinal recording protocols, or can be estimated from the natural-image responses themselves.

## Code and data availability

The complete code to reproduce every experiment and figure in this paper is available at https://github.com/matteofarina2018-cell/CiRF-ViT. The two packaged datasets are available on Zenodo, https://doi.org/10.5281/zenodo.22112881; the underlying recordings were collected by Goldin et al. (2022).

## Acknowledgments

We thank Matthew Chalk for helpful discussions on the LSTA analysis. MF and UF acknowledge this work was done within the framework of the PostGenAI@Paris project with the reference ANR-23-IACL-0007. MF, PZ and UF benefited from financial support by the Agence Nationale de la Recherche (ANR) by the grant IHU FOReSIGHT (ANR-18-IAHU-01). MF and AO received financial support from UKRI AI Centre for Doctoral Training in Biomedical Innovation at the University of Edinburgh. UF thanks Qube Research Technologies (QRT) for their financial support via the project DISSENSATION. MF, PZ and UF’s lab is part of the DIM C-BRAINS, funded by the Conseil Régional d’Île-de-France. PZ’s PhD fellowship was covered by the DIM C-BRAINS, funded by the Conseil Régional d’Île-de-France.

## A Appendix

### A.1 Ablation studies

All ablations follow the same protocol. Each variant changes one element of CiRF-ViT and keeps the rest (tokenizer, readout, loss, optimizer, per-neuron radii and early stopping) unchanged; it is trained five times, once per cross-validation fold, and scored with *CC*_norm_ on the test set (Section A.5). The reference is CiRF-ViT with the five runs at the full training set of Figure 3. In every box plot, each dot is one neuron (its value averaged over the five runs), the box spans the interquartile range, the thick line marks the median and the whiskers the 5th to 95th percentiles; brackets report a two-sided Wilcoxon signed-rank test, paired over neurons (* *p <* 0.05, ** *p <* 0.01, *** *p <* 0.001; ns, not significant). The ablations address, in order, the core and the position of the crop, the components of the model, the radius and the sampling of the crop, and the role of the measured receptive-field centers, in prediction and in the agreement with the LSTA.

#### Attention or convolution, and where the crop acts

The comparison of Figure 3 does not say whether the gain comes from the transformer or from the crop alone, nor whether the crop has to act inside the feature extractor. We therefore trained, with the tokenizer, readout and loss of CiRF-ViT, two cores (Figure 6): the transformer blocks of CiRF-ViT, and convolutional blocks in their place. So that the comparison isolates the operator, the convolutional core matches the transformer blocks in both parameter count and spatial interaction range: each block is *x* + GELU(Conv_1_ × _1_(DWConv_*k*_ × _*k*_(LN(*x*)))), a depthwise *k* × *k* convolution followed by a pointwise one, with *k* = 15 on both datasets, the smallest odd size for which the two blocks connect any two tokens of the median mouse crop, as attention does within the crop in a single layer. The convolutional model has 38,572 parameters on mouse and 46,348 on salamander, against 38,956 and 46,732 for CiRF-ViT. Each core is trained with the crop in two positions. With the *crop in the core*, the blocks only see the tokens inside each neuron’s circle, as in CiRF-ViT (for the convolutional core, the grid outside the circle is held at zero before every convolution). With the *crop in the readout*, the blocks process the whole feature map and the same circle is applied only in the readout, as the exclusion mask of its spatial softmax, so that each neuron’s output pools only the tokens inside its circle.

With the crop in the core, the transformer is better than the convolutional core on both datasets (0.893 against 0.837 on mouse, *p <* 0.001; 0.925 against 0.917 on salamander, *p <* 0.05). On mouse, moving the crop from the readout into the core raises *CC*_norm_ for the transformer, from 0.856 to 0.893 (*p <* 0.01), and for the convolutional core, from 0.812 to 0.837 (*p <* 0.05). On salamander the readout crop is already enough for the transformer (0.927 against 0.925, n.s.), and for the convolutional core the position of the crop makes no difference (0.918 against 0.917, n.s.). A convolutional core that sees as far as attention therefore does not close the gap: the advantage of the transformer is tied to restricting attention itself to each neuron’s receptive field, not to the transformer blocks as such.

#### Component ablations

Figure 7 reports the effect of removing one component of CiRF-ViT at a time, five runs per variant (one per cross-validation fold): the per-neuron crop (every neuron attends to, and is read out from, the whole feature map; the plain ViT of Figure 3), the transformer blocks (tokenizer and readout only), and the self-attention (MLP-only blocks, no token mixing). Each removal lowers *CC*_norm_ on both datasets (*p <* 0.001).

**Figure 7:**
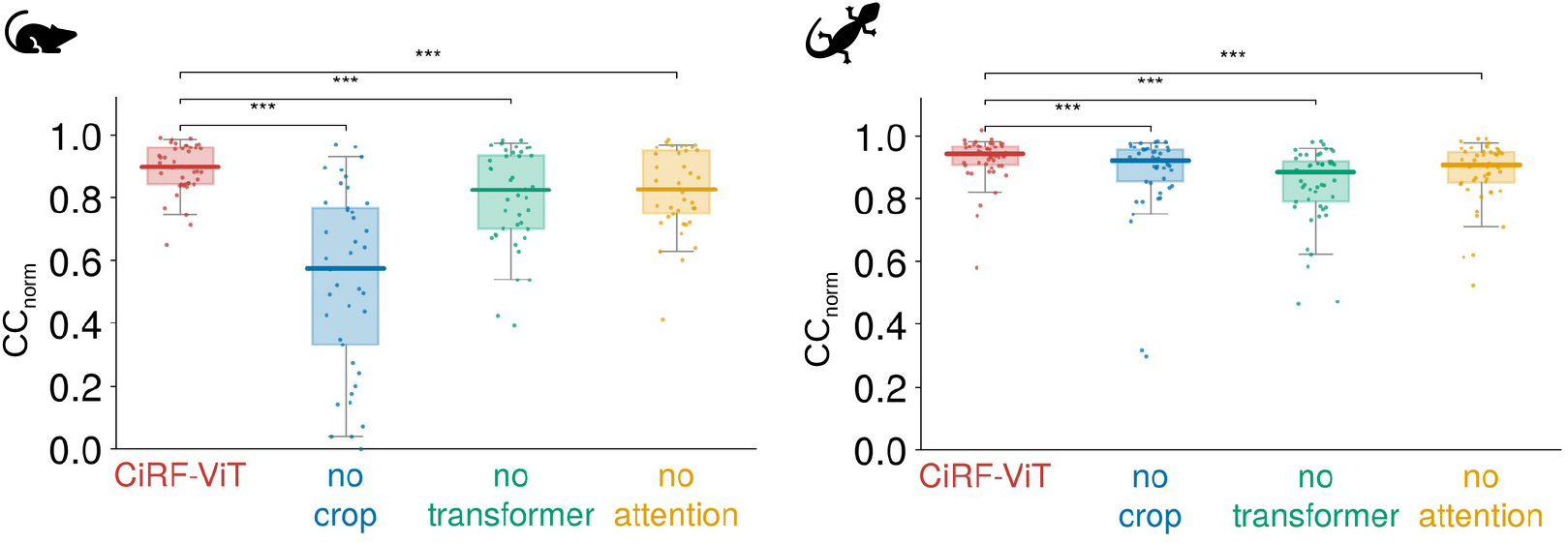
Component ablations. Left: mouse; right: salamander. Per-neuron *CC*_norm_ on the test set, mean of five runs (one per cross-validation fold; for CiRF-ViT, the five runs at the full training set of Figure 3). From left to right: CiRF-ViT; without the per-neuron crop (the plain ViT of Figure 3); without the transformer blocks; without self-attention. Brackets: two-sided Wilcoxon signed-rank test, paired over neurons, between CiRF-ViT and each variant (*** *p <* 0.001). Dots: individual neurons.

#### Crop radius

The crop radii are hyperparameters selected per neuron by Bayesian search. Table 5 scales all of them by the same factor. Shrinking the crop lowers *CC*_norm_ on both datasets (×0.5 and ×0.75, *p <* 0.001; two-sided Wilcoxon signed-rank test, paired over neurons, against CiRF-ViT), whereas enlarging it by half does not lower it significantly, at the cost of more parameters, since the readout grows with the area of the circle. A radius twice as large did not fit in the memory of our GPU.

**Table 5:** Sensitivity to the crop radius: *CC*_norm_ on the test set (mean over five runs ± standard deviation; one run per cross-validation fold, for CiRF-ViT the five runs at the full training set of Figure 3) and number of parameters, with every per-neuron radius multiplied by the factor in the first column.

|  | **Mouse** |  | **Salamander** |  |
|---|---|---|---|---|
| Radius | *CC*<sub>norm</sub> | Parameters | *CC*<sub>norm</sub> | Parameters |
| ×0.5 | 0.845 ± 0.010 | 23,540 | 0.884 ± 0.005 | 28,308 |
| ×0.75 | 0.875 ± 0.008 | 30,920 | 0.905 ± 0.008 | 37,128 |
| ×1 (CiRF-ViT) | 0.893 ± 0.006 | 38,956 | 0.925 ± 0.002 | 46,732 |
| ×1.5 | 0.873 ± 0.010 | 65,196 | 0.920 ± 0.004 | 78,092 |

**Table 6:** Shift of the crop centers during training, from the initial to the trained position (medians over neurons and the five runs). CiRF-ViT starts from the measured receptive-field centers, the random variant of Figure 9 from random positions; *r*_*n*_ is the neuron’s crop radius.

|  | **Mouse** |  | **Salamander** |  |
|---|---|---|---|---|
|  | px | % of *r<sub>n</sub>* | px | % of *r<sub>n</sub>* |
| CiRF-ViT | 0.36 | 3.1 | 0.67 | 4.2 |
| Random centers | 0.34 | 3.1 | 1.07 | 8.0 |

#### Sampling of the crop

Figure 8 replaces the bilinear sampling of the crop (Section A.4) with nearest-neighbor sampling, which is not differentiable with respect to the centers, so that the centers stay at their measured positions, and with bicubic sampling; a fourth variant keeps bilinear sampling but rescales the offsets, so that the tokens lie on a lattice of unit spacing (*s* = 1 instead of *s* = (*W* − 1)*/W*, Section A.4), so that the effective radii equal the values of Table 1 instead of being 1.9% smaller. Bilinear sampling with *s* = (*W* − 1)*/W* gives the highest *CC*_norm_ on both datasets, but the differences are small: at most 0.011 on mouse and 0.020 on salamander.

**Figure 8:**
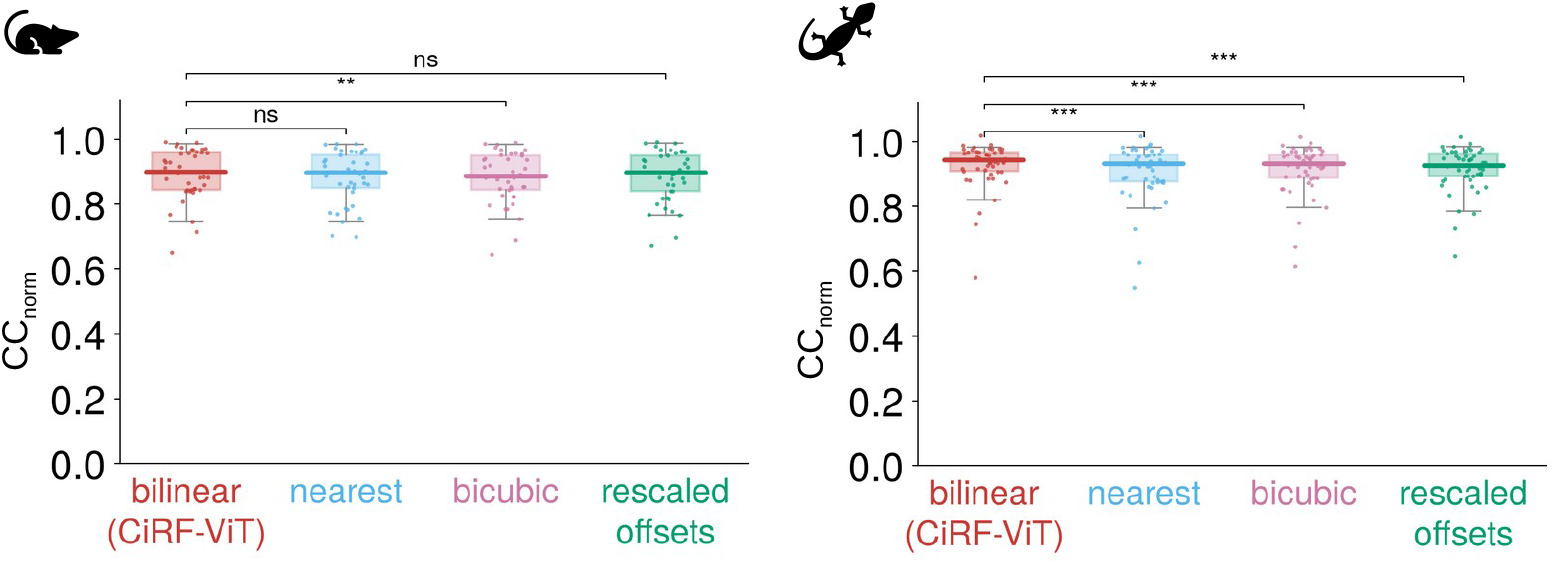
Sampling of the crop. Left: mouse; right: salamander. Per-neuron *CC*_norm_ on the test set, mean of five runs (one per cross-validation fold; for CiRF-ViT, the five runs at the full training set of Figure 3). From left to right: bilinear sampling (CiRF-ViT); nearest-neighbor sampling (centers fixed); bicubic sampling; bilinear sampling with rescaled offsets, on a lattice of unit spacing (*s* = 1, Section A.4). Brackets: two-sided Wilcoxon signed-rank test, paired over neurons, between CiRF-ViT and each variant (** *p <* 0.01, *** *p <* 0.001; ns, not significant). Dots: individual neurons.

#### Initialization of the crop centers

CiRF-ViT initializes each neuron’s crop center at the peak of its STA, measured in a separate white-noise experiment. Figure 9 replaces this initialization with four alternatives that use less or none of that measurement, with the per-neuron radii unchanged and the centers trained in every case: the centroid of the neuron’s cluster of measured centers (*k*-means, *k* = 4); a single center shared by all neurons, the population mean of the measured centers; centers estimated from the natural-image training set alone, without the STA, as the peak of the smoothed absolute correlation between the neuron’s response and each pixel; and centers drawn uniformly over the image.

The model does not need the precision of the individual measurement: with centers known only up to their cluster, *CC*_norm_ is 0.885 on mouse (against 0.893, n.s.) and 0.916 on salamander (against 0.925), and a single shared center still reaches 0.844 and 0.902. Centers estimated from the training data, without any external measurement, reproduce most of the result: the median neuron is predicted almost as well as by CiRF-ViT (0.868 against 0.899 on mouse, 0.925 against 0.943 on salamander), and 73% and 80% of the neurons lose less than 0.05. The loss of the mean (0.766 and 0.890) comes from a few neurons whose estimated center is far from the receptive field (6 of 41 on mouse and 1 of 49 on salamander lose more than 0.3). With random centers the model fails (0.233 and 0.416): starting far from the receptive field, the centers do not reach it during training.

#### Displacement of the crop centers

The crop centers move little during training, wherever they start: the median shift is a fraction of a pixel to about a pixel, a few percent of the crop radius, both for CiRF-ViT, which starts from the measured centers, and for the random initialization of Figure 9 (Table 6). Random centers, which start about 40 pixels from the measured center, therefore do not reach the receptive field: their distance to it is essentially unchanged after training (Figure 10), which explains the failure of the random initialization in Figure 9.

**Figure 9:**
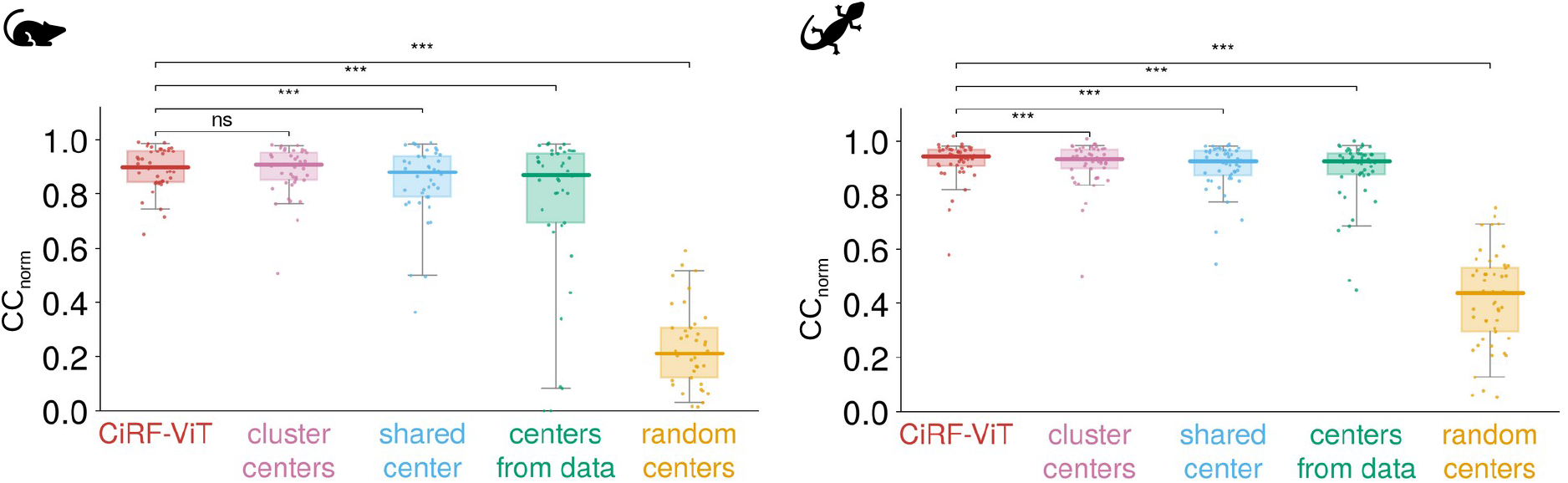
Initialization of the crop centers. Left: mouse; right: salamander. Per-neuron *CC*_norm_ on the test set, mean of five runs (one per cross-validation fold; for CiRF-ViT, the five runs at the full training set of Figure 3). From left to right: CiRF-ViT (centers initialized at the measured receptive-field centers); centroid of the neuron’s cluster of measured centers; one center shared by all neurons; centers estimated from the training responses alone; random centers. Brackets: two-sided Wilcoxon signed-rank test, paired over neurons, between CiRF-ViT and each variant (*** *p <* 0.001; ns, not significant). Dots: individual neurons.

**Figure 10:**
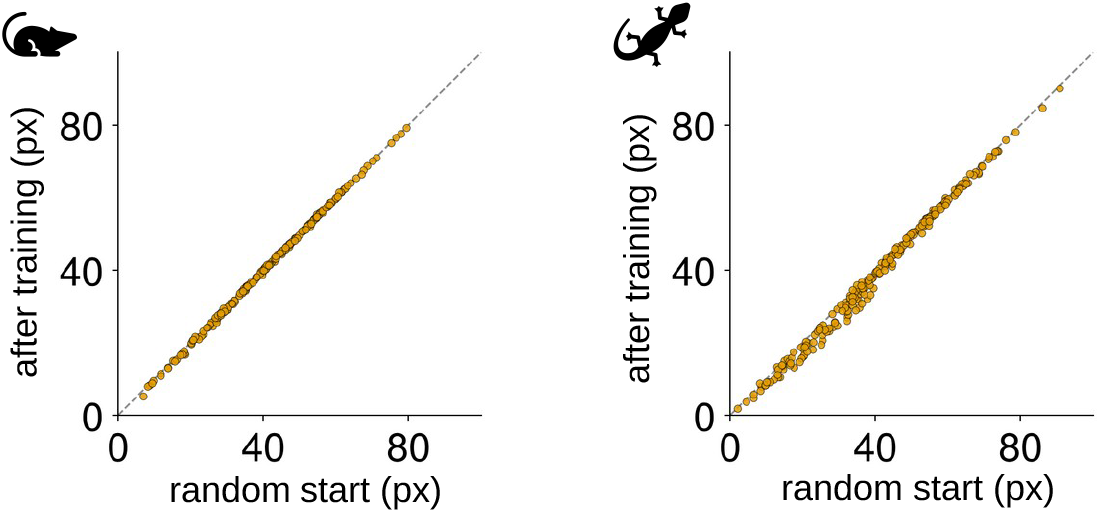
Random crop centers do not migrate to the receptive field. Left: mouse; right: salamander. Distance between the crop center and the measured receptive-field center (STA peak), in stimulus pixels, at the random initialization (x-axis) and after training (y-axis), for the random-center variant of Figure 9. Each dot is one neuron in one of the five runs; the dashed line is the identity.

#### LSTA agreement without the measured receptive field

CiRF-ViT receives, of the STA, only the location of its peak, used to initialize the crop centers. Figure 11 repeats the LSTA comparison of Section 6 for models that never receive the STA. With centers estimated from the natural-image training responses (Figure 9), the agreement with the experimental LSTA is somewhat lower than that of CiRF-ViT (*p <* 0.001; median over neurons 0.519 against 0.567 on mouse, 0.445 against 0.503 on salamander), as is the prediction. The plain ViT, which has no crop and no center at all, matches CiRF-ViT on salamander (n.s.), where its prediction is only moderately lower (*CC*_norm_ 0.88 against 0.93), and fails on mouse, where its prediction fails too (0.54 against 0.89; Figure 3). With random centers, the agreement vanishes on both datasets, as does the prediction. The agreement with the LSTA therefore follows the quality of the prediction, not the information the model receives about the STA.

**Figure 11:**
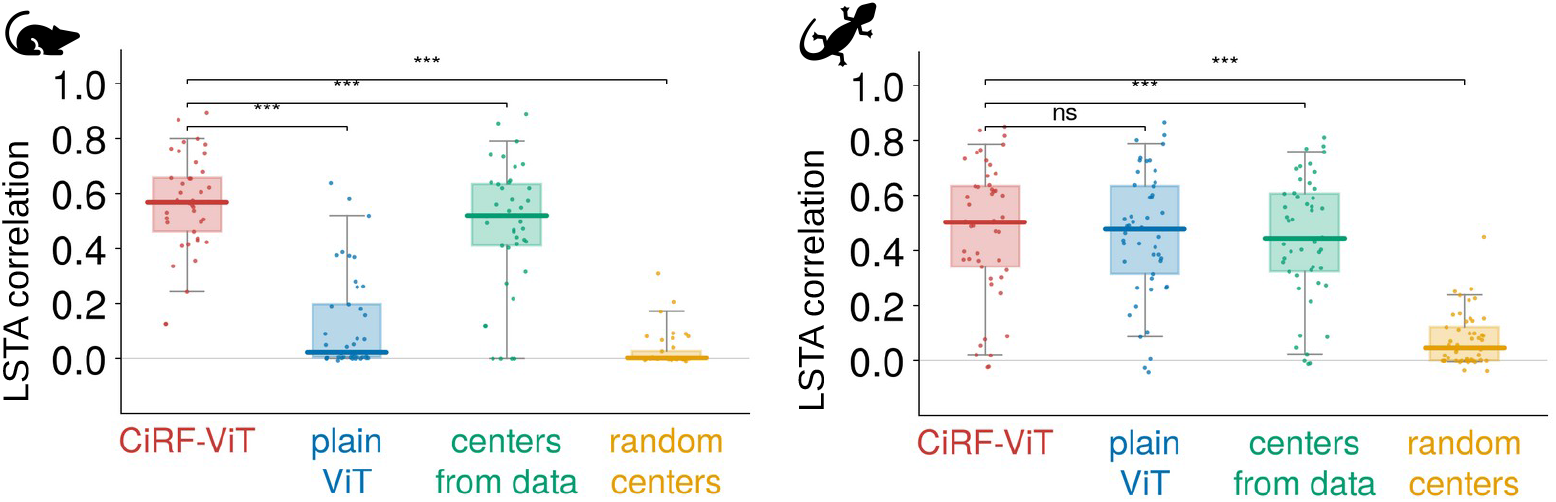
Agreement with the experimental LSTA of models that do not receive the STA. Left: mouse; right: salamander. Per-neuron correlation between the experimental LSTA and the model gradient, band-limited as in Figure 4e, mean of five runs (one per cross-validation fold; for CiRF-ViT, the five runs at the full training set of Figure 3). From left to right: CiRF-ViT; plain ViT; centers estimated from the training responses; random centers. Brackets: two-sided Wilcoxon signed-rank test, paired over neurons, between CiRF-ViT and each variant (*** *p <* 0.001; ns, not significant). Dots: individual neurons.

### A.2 Pretrained transformer cores

An alternative to a locality prior, when data are scarce, is to start from a transformer pretrained on other data. We replaced the tokenizer and transformer blocks of CiRF-ViT with three pretrained cores, each followed by the readout of CiRF-ViT over the whole token grid (without the crop) and trained with the same loss, partition and early stopping: a ViT-S/16 supervised on ImageNet (Dosovitskiy et al., 2021; Steiner et al., 2022), a ViT-S/16 pretrained with DINO on ImageNet (Caron et al., 2021), and the V1T core of Li et al. (2023), trained on the responses of mouse V1 to natural images. Each core is either frozen, with only the readout trained, or fully fine-tuned. For the two ImageNet models, the number of transformer blocks kept was chosen on the validation loss (one to three blocks, except for the fine-tuned DINO model on salamander, which keeps all twelve). The stimulus is resized to the input of each model (224 × 224 pixels on three channels for the ImageNet models, 36 × 36 for V1T).

None of the pretrained cores reaches CiRF-ViT at any training-set size (Figure 12). The best is V1T, the only one pretrained on neural responses: with the full training set, fine-tuned, it reaches 0.69 on mouse and 0.83 on salamander, against 0.89 and 0.93 for CiRF-ViT and 0.87 and 0.86 for the CNN. Its gap to CiRF-ViT is smallest with 10% of the data (0.50 against 0.56 on mouse, 0.63 against 0.71 on salamander); on mouse, with 10 and 20% of the data, it is the only pretrained core above the CNN.

**Figure 12:**
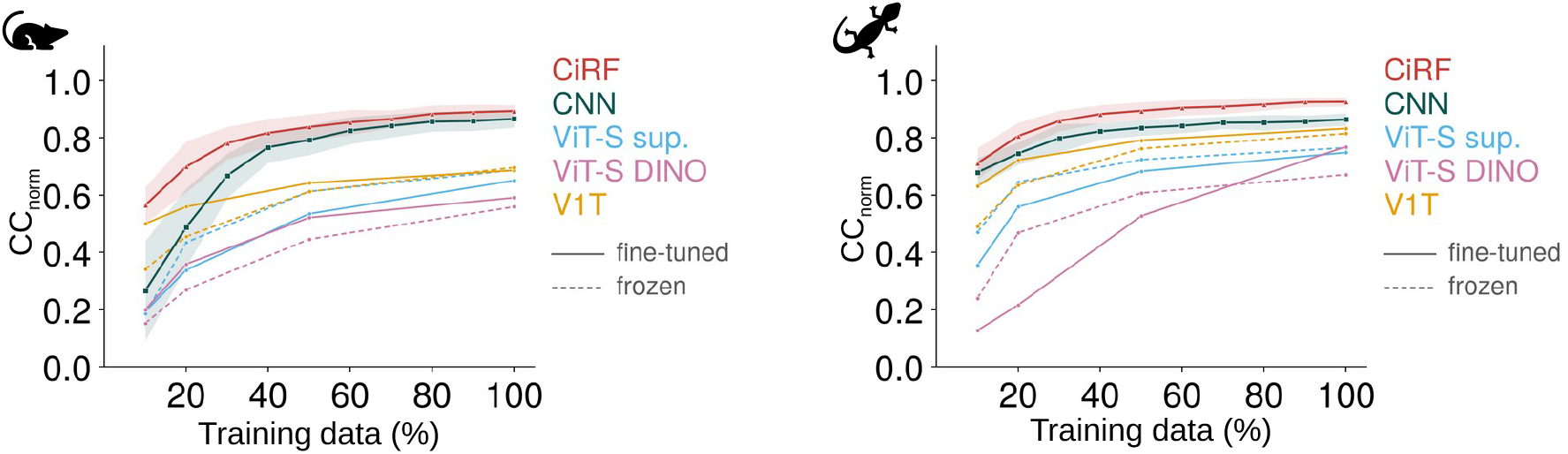
Pretrained transformer cores. Left: mouse; right: salamander. *CC*_norm_ (mean over neurons, averaged over runs) as a function of the fraction of training data, as in Figure 3c. CiRF-ViT and the CNN are the curves of Figure 3c (five runs per point; shaded regions as there). Pretrained cores, with the readout of CiRF-ViT over the whole token grid: ViT-S/16 supervised on ImageNet (ViT-S sup.), ViT-S/16 pretrained with DINO (ViT-S DINO), and the V1T core of Li et al. (2023) (V1T); solid lines, fully fine-tuned; dashed lines, frozen core. Five runs per point (one per cross-validation fold), at 10, 20, 50 and 100% of the training data.

### A.3 Training cost

Table 7 reports the training cost of the models of Figure 6 on the mouse dataset, on one NVIDIA GB10 GPU; for the convolutional core, it reports an earlier version with 3 × 3 convolutions in place of the 15 × 15 depthwise convolutions of Figure 6, which were not timed. The time per step covers the forward pass, the Poisson loss, the backward pass and the Adam update on one batch of 32 images. The transformer models run without FlashAttention, whose kernels are not available on this GPU.

**Table 7:**
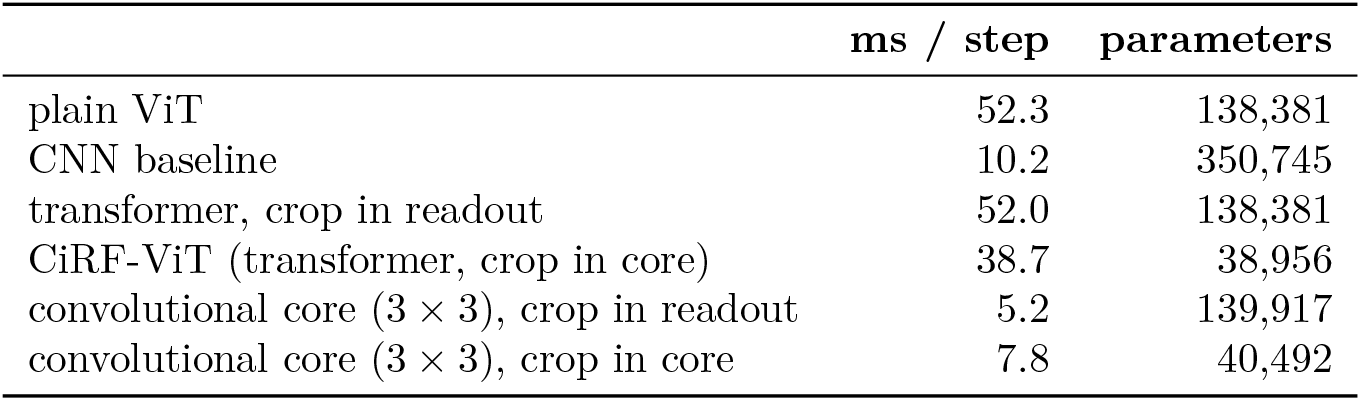
Training cost on the mouse dataset.

**Table 8:**
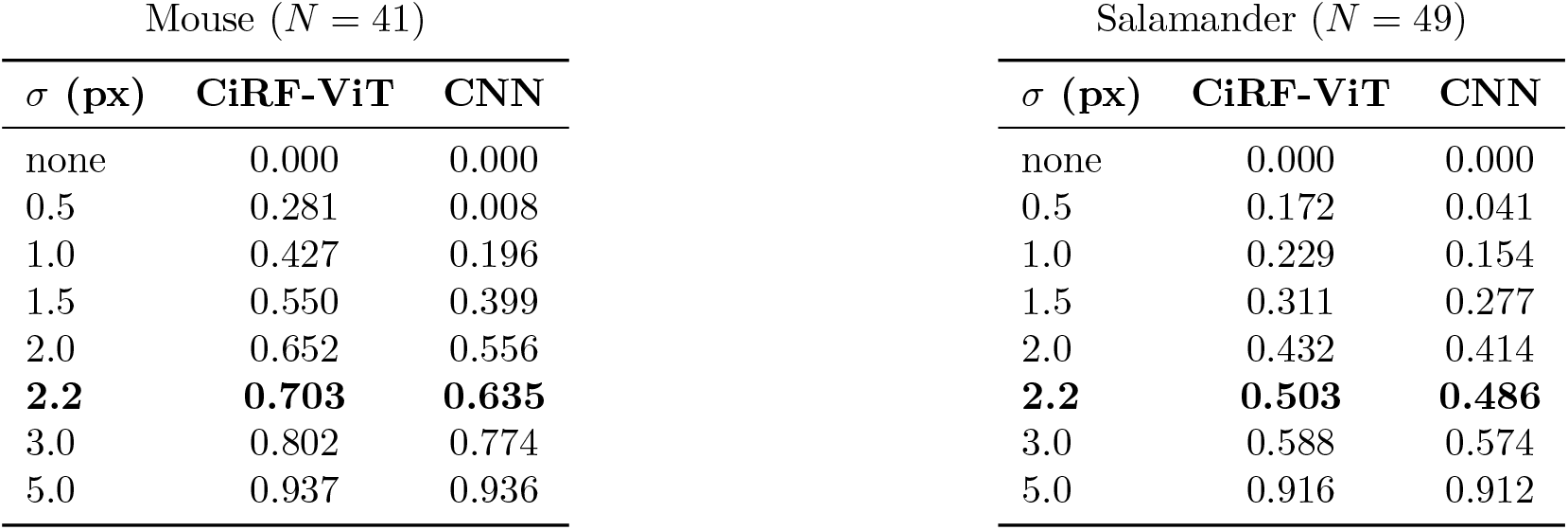
Fraction of input-gradient power that a Gaussian low-pass of width *σ* removes, 1 −∥*G*_*σ*_ \**g*∥ ^2^*/* ∥*g*∥ ^2^ for a gradient map *g* and the Gaussian kernel *G*_*σ*_. A Gaussian filter has no sharp edge, so this is an attenuation weighted over all frequencies, not the power strictly above a cutoff; the attenuation is concentrated above the half-amplitude frequency *f*_*c*_ = 0.187*/σ* cycles per pixel, that is, on spatial scales finer than a period of about 5.34 *σ* px. Measured inside the receptive-field window on the 108 × 108 stimulus frame, averaged over ten independently trained models of each architecture, all neurons and all test images. The first row applies no cut and is 0 by definition; the fraction grows with *σ* because a wider kernel moves the attenuation to lower frequencies and so attenuates more of the band. The row in bold, *σ* = 2.2, is the value used to band-limit the maps compared with the experimental LSTA in Figure 4.

| Mouse (*N* = 41) |  |  | Salamander (*N* = 49) |  |  |
|---|---|---|---|---|---|
| *σ* **(px)** | **CiRF-ViT** | **CNN** | *σ* **(px)** | **CiRF-ViT** | **CNN** |
| none | 0.000 | 0.000 | none | 0.000 | 0.000 |
| 0.5 | 0.281 | 0.008 | 0.5 | 0.172 | 0.041 |
| 1.0 | 0.427 | 0.196 | 1.0 | 0.229 | 0.154 |
| 1.5 | 0.550 | 0.399 | 1.5 | 0.311 | 0.277 |
| 2.0 | 0.652 | 0.556 | 2.0 | 0.432 | 0.414 |
| **2.2** | **0.703** | **0.635** | **2.2** | **0.503** | **0.486** |
| 3.0 | 0.802 | 0.774 | 3.0 | 0.588 | 0.574 |
| 5.0 | 0.937 | 0.936 | 5.0 | 0.916 | 0.912 |

The crop also reduces the number of tokens each neuron attends to. Global attention sees the whole 54 × 54 feature map, 2916 tokens, for every neuron; with the crop, a neuron attends to the tokens inside its circle, 137 for the median neuron on mouse and 177 on salamander (between 13 and 489, depending on the radius), that is 4.7% and 6.1% of the map. The transformer blocks are shared and do not depend on the number of tokens, so the parameter counts of CiRF-ViT and of the plain ViT differ only in the readout, whose spatial weights cover the circle instead of the whole map, and in the two trainable center coordinates of each neuron (Table 7).

These counts describe what each neuron can attend to, not what the implementation computes. To keep tensor shapes static, all neurons share one set of *T*_circle_ = 489 tokens, fixed by the largest radius (12.5 feature-map positions on both datasets), and the tokens outside each neuron’s circle are excluded by a key-padding mask. This length is set by the few neurons with the largest radius (4 of 41 on mouse and 3 of 49 on salamander) and is much larger than most neurons need: the median neuron uses 137 of the 489 tokens on mouse and 177 on salamander. Dense attention still computes the scores of the masked tokens, so every neuron costs 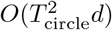 whatever its radius, and the transformer runs once per neuron: 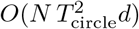 per image and layer, against *O*((*HW*)^2^*d*) once per image for the plain ViT, whose output is the same for all neurons. Per image, layer and head, this is 9.8M attention scores on mouse (*N* = 41) and 11.7M on salamander (*N* = 49), against 8.5M for the plain ViT; with variable-length attention, each neuron restricted to its own tokens, it would be 2.2M and 3.4M, so 77% of the scores computed on mouse and 71% on salamander are discarded by the mask. The token-wise layers (projections and MLP) process *N T*_circle_ = 20,049 and 23,961 tokens, against 2,916. The theoretical saving of the crop is therefore not realized by our implementation, whose attention costs about as much as global attention; realizing it would need variable-length or block-sparse attention kernels. The measured time per step of CiRF-ViT is nevertheless lower than that of the plain ViT (Table 7). The advantage of the crop reported in this work is therefore not in training time but in how well the model learns from few data: with the same training set and less than a third of the parameters, CiRF-ViT reaches a *CC*_norm_ of 0.893 on mouse and 0.925 on salamander, against 0.542 and 0.882 for the plain ViT, and it stays competitive with the GP baseline when the training set is reduced to 10–20% of its size (Figure 3c).

### A.4 Differentiable per-neuron circular crop

Let *F* ∈ ℝ^*C×H×W*^ be the feature map produced by the CNN tokenizer for one input image, where *C* is the channel dimension and (*H, W*) its spatial resolution, and let *N* be the number of neurons. Throughout this section, positions on the feature map are written as (row, column) pairs, that is, (*y, x*), and *F* [:, *i, j*] denotes the feature vector at row *i* (along *y*) and column *j* (along *x*). Each neuron *n* is associated with a center 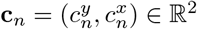 and a radius *r*_*n*_ *>* 0, both expressed in feature-map pixel coordinates (the radii of Table 1, given in image pixels, divided by the tokenizer stride). The centers are initialized at the receptive-field centers estimated from the spike-triggered average (Section 3), mapped to feature-map coordinates, and stored as free parameters optimized by backpropagation jointly with all other model parameters; the radii are instead fixed hyperparameters, selected per neuron by Bayesian hyperparameter search and kept constant during training. Since the radii differ across neurons, we define a single shared set of sampling offsets on the integer lattice *G* = {−*h*, …, *h*} ^2^, with *h* = ⌈*r*_max_⌉ and *r*_max_ = max_*n*_ *r*_*n*_, and retain the offsets contained in the largest circle,

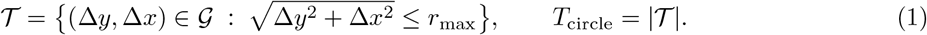

In this way the number of tokens *T*_circle_ is identical for every neuron and every batch, keeping the computation statically shaped, and since the offsets are integer-valued the set *T* is precomputed once at initialization. The specificity of each neuron within this shared token set is carried by a static binary mask,

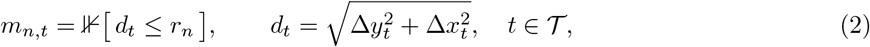

whose complement is applied as a key-padding mask in the self-attention, by setting the attention logits over excluded tokens to −∞ before the softmax, and as an exclusion mask in the spatial softmax of the readout, so that tokens outside *r*_*n*_ contribute exactly zero to the neuron’s output. The crop itself is implemented as differentiable resampling: for neuron *n* and offset *t*, the sampling location **p**_*n,t*_ = **c**_*n*_ + *s* (Δ*y*_*t*_, Δ*x*_*t*_), with the lattice spacing *s* = (*W* − 1)*/W* defined below, generally falls between pixel centers because **c**_*n*_ is continuous, and the corresponding feature vector is obtained by bilinear interpolation of the four neighboring pixels. Writing **p**_*n,t*_ = (*p*^*y*^, *p*^*x*^), *i*_0_ = ⌊*p*^*y*^⌋, *j*_0_ = ⌊*p*^*x*^⌋, and *v* = *p*^*y*^ − *i*_0_, *u* = *p*^*x*^ − *j*_0_ for the fractional offsets along *y* and *x*,

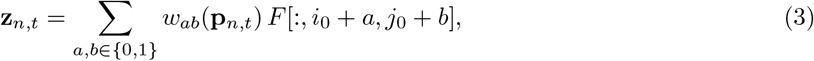

where the first index of the weights refers to the row (*y*) and the second to the column (*x*): *w*_00_ = (1−*v*)(1−*u*), *w*_01_ = (1 − *v*)*u, w*_10_ = *v*(1 − *u*), *w*_11_ = *vu*. The weights have piecewise-constant derivatives with respect to the center coordinates, e.g. 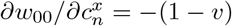, since 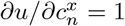. Gradients of the loss therefore flow to 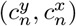 through the interpolation weights, making the center of each neuron’s crop trainable while the offsets in remain fixed. In practice the sampling uses `grid_sample` in bilinear mode with border padding and `align_corners=True`, under which the normalized coordinates − 1 and +1 refer to the centers of the first and last pixel. The centers are mapped accordingly, 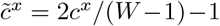 and 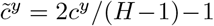, and the offsets as 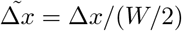 and 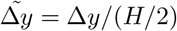, which gives exactly the sampling location **p**_*n,t*_ = **c**_*n*_ +*s* (Δ*y*_*t*_, Δ*x*_*t*_) with *s* = (*W* − 1)*/W* = 53*/*54 ≈ 0.981 on the 54 × 54 feature map. The tokens therefore lie on a lattice slightly finer than the pixel grid, and the effective radius of neuron *n* is *s r*_*n*_, with *r*_*n*_ the value in Table 1. We kept this spacing rather than *s* = 1 because the radii were selected by Bayesian search, and every result in this paper was obtained with it: changing *s* changes the effective radius of every neuron and the sampling lattice, and a fair comparison would require repeating the search and every experiment. With *s* = 1 and the same radii the model is not better (Figure 8: 0.885 against 0.893 on mouse, 0.914 against 0.925 on salamander), and the released code provides both spacings. The output of this stage is, for each image, a tensor of tokens **z**_*n,t*_ ∈ ℝ^*C*^ of shape [*N, T*_circle_, *C*], together with the per-neuron masks *m*_*n,t*_. Because token *t* is sampled at the same offset (Δ*y*_*t*_, Δ*x*_*t*_) from every neuron’s center, the token index itself defines the position relative to the neuron, and the token representation is invariant to the center location. Tokens and masks are then fed to the shared-weight transformer blocks described in Section 4.2.

### A.5 Definition of the prediction metric

For each neuron, the repetitions of the test responses are split into two halves (even- and odd-numbered trials), and each half is averaged into a response vector over the 30 test images, 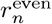 and 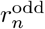. Writing for the Pearson correlation across images and *ŷ*_*n*_ for the predicted rate, the per-neuron score is

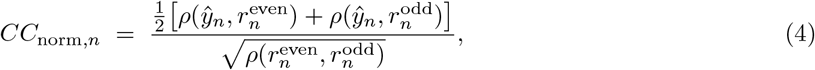

clipped below at zero, and set to zero when the half-to-half correlation is not positive. This is the noise-corrected correlation of Schoppe et al. (2016), with the ceiling estimated from the two halves. If each half is the noise-free response plus independent repetition noise, the correlation between the halves is the product of the correlations of the noise-free response with each half, 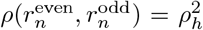; a model that predicts the noise-free response correlates with each half by *ρ*_*h*_ and therefore scores 1, which is why the denominator is a square root. Its square is the fraction of the explainable variance captured by the prediction (the adjusted *R*^2^). Individual values can still exceed 1 through the sampling error of the correlations; the axes of Figure 3 extend accordingly. Each per-neuron value rests on correlations over 30 images and thus carries a non-negligible sampling error; the population summaries and the paired tests, not the individual points, are the quantities the figures are meant to convey.

### A.6 Band-limiting the LSTA comparison

The quantitative comparison has to respect what the experiment can measure. The correlation of Figure 4e compares a model’s input gradient with the experimental LSTA inside the receptive-field window. The LSTA is estimated from a finite number of spikes, and at fine spatial scales the estimate is dominated by noise: a spectral analysis of the experimental maps, contrasting each neuron’s receptive-field window with an identical window placed far from the receptive field, shows that above a cutoff frequency the two are indistinguishable, so that band carries no signal for a model to agree with. Both the experimental map and the model gradients nevertheless carry energy at spatial frequencies where the experiment measures only noise. That energy cannot contribute to the covariance between the two maps, but it does enter the model’s variance in the denominator of the correlation, so an unfiltered correlation penalizes a model for having any content in a band where there is nothing to agree with; the penalty is not neutral across models, since the rougher a model’s gradients, the larger it is. We therefore low-pass both sides with the same Gaussian filter before correlating, removing the noise-dominated band from the comparison. This is a change to the measurement, not to the models: it does not assert that the suppressed structure is noise on the model side, only that the experiment cannot be used to check it.

#### Choosing the filter width

The cutoff is derived from the experimental data. For each neuron we take two windows of identical size at the LSTA’s native resolution (72 px frames on mouse, 56 px on salamander): a signal window, the bounding box of the receptive-field ellipse plus the same padding the correlation uses, and a noise window, the same box pushed to the opposite side of the frame, far from the receptive field, where the neuron does not respond and the map contains estimation noise only. Spectra are estimated at the native resolution, because estimating them after resampling to the 108 px analysis frame would fold the interpolation’s own low-pass into the answer. Averaging Hann-windowed radial power spectra over all neurons and images (328 windows on mouse, 196 on salamander) gives a per-frequency signal-to-noise ratio, and the usable band is defined as the frequencies where it exceeds 2. The SNR crosses that threshold at a spatial period of 7.4 px native on mouse and 6.4 px on salamander, that is, 11.1 and 12.3 px once expressed in the 108 × 108 frame.

A Gaussian low-pass of standard deviation *σ* has transfer function exp(−2*π*^2^*σ*^2^*f* ^2^) for spatial frequency *f* in cycles per pixel, with half-amplitude point, where the transfer function falls to one half (−6 dB), at *f*_*c*_ = 0.187*/σ*, a spatial period of 5.34 *σ* px; the half-power point (−3 dB) lies lower, at 0.132*/σ*. Placing the half-amplitude point at the measured cutoff gives *σ* = 2.08 px for mouse and *σ* = 2.31 px for salamander; we use their mean, *σ* = 2.2 px in the 108 × 108 frame, as a common value for both datasets. The filter is applied identically to the experimental LSTA and to every model’s gradients, after resampling to the analysis frame. Before the correlation, both maps then undergo the same post-processing: each map is squared with its sign preserved, and pixels below 20% of the map’s peak absolute value are set to zero. Both steps suppress the low-amplitude background noise that survives the filter, and are applied identically to both sides for every model.

#### What the filter removes from each model

Table 8 reports the fraction of input-gradient power that a low-pass of width *σ* removes (plotted in Figure 4f), for CiRF-ViT and the CNN, averaged over ten independently trained models of each architecture; the GP is omitted because its kernel makes its gradients smooth by construction, so the informative contrast is between the two gradient-trained architectures. At the working value *σ* = 2.2 (bold row) the filter removes 70% of CiRF-ViT’s gradient power on mouse and 50% on salamander, against 64% and 49% for the CNN. The difference is concentrated at the finest scales: a filter of width *σ* = 0.5, whose attenuation is concentrated on periods finer than about 2.7 px, strips 28% of CiRF-ViT’s power on mouse but less than 1% of the CNN’s, and 17% against 4% on salamander. CiRF-ViT is therefore far richer in high-frequency components than the CNN, and it is exactly this content that the band limit excludes from Figure 4e: it lies above the frequencies the experimental LSTA resolves, so these data can neither confirm nor refute it.

## References

Santiago A. Cadena, George H. Denfield, Edgar Y. Walker, Leon A. Gatys, Andreas S. Tolias, Matthias Bethge, and Alexander S. Ecker. Deep convolutional models improve predictions of macaque V1 responses to natural images. PLOS Computational Biology, 15(4):e1006897, 2019.

Mathilde Caron, Hugo Touvron, Ishan Misra, Hervé Jégou, Julien Mairal, Piotr Bojanowski, and Armand Joulin. Emerging properties in self-supervised vision transformers. In Proceedings of the IEEE/CVF International Conference on Computer Vision, pp. 9650–9660, 2021.

Bin Chen, Ran Wang, Di Ming, and Xin Feng. ViT-P: Rethinking data-efficient vision transformers from locality. arXiv preprint arXiv:2203.02358, 2022.

Youngmin Cho and Lawrence K. Saul. Kernel methods for deep learning. In Advances in Neural Information Processing Systems, volume 22, pp. 342–350, 2009.

Jean-Baptiste Cordonnier, Andreas Loukas, and Martin Jaggi. On the relationship between self-attention and convolutional layers. In International Conference on Learning Representations, 2020.

Stéphane d’Ascoli, Hugo Touvron, Matthew L. Leavitt, Ari S. Morcos, Giulio Biroli, and Levent Sagun. Con-ViT: Improving vision transformers with soft convolutional inductive biases. In International Conference on Machine Learning, 2021.

Alexey Dosovitskiy, Lucas Beyer, Alexander Kolesnikov, Dirk Weissenborn, Xiaohua Zhai, Thomas Unterthiner, Mostafa Dehghani, Matthias Minderer, Georg Heigold, Sylvain Gelly, Jakob Uszkoreit, and Neil Houlsby. An image is worth 16x16 words: Transformers for image recognition at scale. In International Conference on Learning Representations, 2021.

Matías A. Goldin, Baptiste Lefebvre, Samuele Virgili, Mathieu Kim Pham Van Cang, Alexander Ecker, Thierry Mora, Ulisse Ferrari, and Olivier Marre. Context-dependent selectivity to natural images in the retina. Nature Communications, 13(1):5556, 2022. doi: 10.1038/s41467-022-33242-8.

Matías A. Goldin, Samuele Virgili, and Matthew Chalk. Scalable Gaussian process inference of neural responses to natural images. Proceedings of the National Academy of Sciences, 120(34):e2301150120, 2023. doi: 10.1073/pnas.2301150120.

Bum Jun Kim, Hyeyeon Choi, Hyeonah Jang, Dong Gu Lee, Wonseok Jeong, and Sang Woo Kim. Dead pixel test using effective receptive field. Pattern Recognition Letters, 167:149–156, 2023.

David A. Klindt, Alexander S. Ecker, Thomas Euler, and Matthias Bethge. Neural system identification for large populations separating “what” and “where”. In Advances in Neural Information Processing Systems, volume 30, pp. 3509–3519, 2017.

Bryan M. Li, Isabel M. Cornacchia, Nathalie L. Rochefort, and Arno Onken. V1T: Large-scale mouse V1 response prediction using a vision transformer. Transactions on Machine Learning Research, 2023. URL https://openreview.net/forum?id=qHZs2p4ZD4.

Ze Liu, Yutong Lin, Yue Cao, Han Hu, Yixuan Wei, Zheng Zhang, Stephen Lin, and Baining Guo. Swin Transformer: Hierarchical vision transformer using shifted windows. In International Conference on Computer Vision, 2021.

Wenjie Luo, Yujia Li, Raquel Urtasun, and Richard Zemel. Understanding the effective receptive field in deep convolutional neural networks. In Advances in Neural Information Processing Systems, volume 29, 2016.

Konstantin-Klemens Lurz, Mohammad Bashiri, Konstantin Willeke, Akshay K. Jagadish, Eric Wang, Edgar Y. Walker, Santiago A. Cadena, Taliah Muhammad, Erick Cobos, Andreas S. Tolias, Alexander S. Ecker, and Fabian H. Sinz. Generalization in data-driven models of primary visual cortex. In International Conference on Learning Representations, 2021.

Oliver Schoppe, Nicol S. Harper, Ben D. B. Willmore, Andrew J. King, and Jan W. H. Schnupp. Measuring the performance of neural models. Frontiers in Computational Neuroscience, 10:10, 2016. doi: 10.3389/fncom.2016.00010.

Andreas Steiner, Alexander Kolesnikov, Xiaohua Zhai, Ross Wightman, Jakob Uszkoreit, and Lucas Beyer. How to train your ViT? Data, augmentation, and regularization in vision transformers. Transactions on Machine Learning Research, 2022.

Hugo Touvron, Matthieu Cord, Matthijs Douze, Francisco Massa, Alexandre Sablayrolles, and Hervé Jégou. Training data-efficient image transformers & distillation through attention. In International Conference on Machine Learning, pp. 10347–10357, 2021.

Frank Wilcoxon. Individual comparisons by ranking methods. Biometrics Bulletin, 1(6):80–83, 1945. doi: 10.2307/3001968.

Tete Xiao, Mannat Singh, Eric Mintun, Trevor Darrell, Piotr Dollár, and Ross Girshick. Early convolutions help transformers see better. In Advances in Neural Information Processing Systems, 2021.

